# Vaccinia virus-based vaccines confer protective immunity against SARS-CoV-2 virus in Syrian hamsters

**DOI:** 10.1101/2021.08.03.454910

**Authors:** Rakesh Kulkarni, Wen-Ching Chen, Ying Lee, Chi-Fei Kao, Shiu-Lok Hu, Hsiu-Hua Ma, Jia-Tsrong Jan, Chun-Che Liao, Jian-Jong Liang, Hui-Ying Ko, Cheng-Pu Sun, Yin-Shoiou Lin, Yu-Chiuan Wang, Sung-Chan Wei, Yi-Ling Lin, Che Ma, Yu-Chan Chao, Yu-Chi Chou, Wen Chang

**Author notes:** Corresponding author (WC).

## Abstract

COVID-19 in humans is caused by Severe acute respiratory syndrome coronavirus-2 (SARS-CoV-2) that belongs to the beta family of coronaviruses. SARS-CoV-2 causes severe respiratory illness in 10-15% of infected individuals and mortality in 2-3%. Vaccines are urgently needed to prevent infection and to contain viral spread. Although several mRNA- and adenovirus-based vaccines are highly effective, their dependence on the “cold chain” transportation makes global vaccination a difficult task. In this context, a stable lyophilized vaccine may present certain advantages. Accordingly, establishing additional vaccine platforms remains vital to tackle SARS-CoV-2 and any future variants that may arise. Vaccinia virus (VACV) has been used to eradicate smallpox disease, and several attenuated viral strains with enhanced safety for human applications have been developed. We have generated two candidate SARS-CoV-2 vaccines based on two vaccinia viral strains, MVA and v-NY, that express full-length SARS-CoV-2 spike protein. Whereas MVA is growth-restricted in mammalian cells, the v-NY strain is replication-competent. We demonstrate that both candidate recombinant vaccines induce high titers of neutralizing antibodies in C57BL/6 mice vaccinated according to prime-boost regimens. Furthermore, our vaccination regimens generated T_H_1-biased immune responses in mice. Most importantly, prime-boost vaccination of a Syrian hamster infection model with MVA-S and v-NY-S protected the hamsters against SARS-CoV-2 infection, supporting that these two vaccines are promising candidates for future development. Finally, our vaccination regimens generated neutralizing antibodies that partially cross-neutralized SARS-CoV-2 variants of concern.

## Introduction

Severe acute respiratory syndrome coronavirus 2 (SARS-CoV-2), a member of the *Betacoronavirus* family, is causing a global pandemic and, as of July 2021, has infected more than 190 million people worldwide and resulted in 4 million deaths (https://covid19.who.int/) (1, 2). Compared to two other highly pathogenic coronaviruses, SARS-CoV (3) and Middle east respiratory syndrome coronavirus (MERS-CoV) (4), SARS-CoV-2 has proven more difficult to contain (5). Consequently, an effective vaccine to halt the spread of SARS-CoV-2 is urgently needed.

SARS-CoV-2 is an enveloped single-stranded positive-sense RNA virus, whose Spike protein (S) on the virion surface mediates virus entry into target cells (6–8). Spike protein has S1 and S2 components and, similar to other type 1 viral fusion proteins, the S1 subunit contains a receptor-binding domain (RBD) that binds to its host cell receptor, angiotensin converting enzyme 2 (ACE2) (9), whereas the S2 subunit mediates membrane fusion (10). The S protein of some SARS-CoV-2 strains requires cleavage by the cellular serine protease TMPRSS2 during cell entry (8, 11). Neutralizing antibodies from convalescent patients recognize S protein, making it a good vaccine target (12, 13). S protein is also a major target of T cell responses to SARS-CoV-2 (14, 15). Although several SARS-CoV-2 vaccines, developed using mRNA technology (16–18) and adenovirus vectors (19–21), are currently in use; however, additional vaccines that are cost effective and could be transported without cold chain will still be worthwhile to develop. In addition, concerns have been raised of adverse effects following vaccination (22–24), implying that improvements to currently available SARS-CoV-2 vaccines are essential and will necessitate ongoing vaccine development.

Vaccinia virus has been deployed successfully to eradicate smallpox worldwide (25, 26). The Modified Vaccinia Ankara (MVA) strain is growth-restricted in mammalian cells and preclinical and clinical trials have demonstrated it to be quite a safe vaccine vector against viral diseases such as HIV, MERS-CoV and SARS-CoV (27–30). However, other attenuated strains of vaccinia virus exhibiting different degrees of immunogenicity could also serve as vaccine vectors (31–40). Recently, several reports revealed that the MVA strain expressing SARS-CoV-2 S protein protected ACE2-transgenic mice and macaques from SARS-CoV-2 challenges (41–43). Here, we generated SARS-CoV-2 vaccines using the MVA strain, as well as a v- NY strain previously employed as a vector for the first recombinant vaccinia virus (HIVAC-1e) used in FDA-approved clinical trials (44–48), both of which we engineered to express SARS-CoV-2 S protein. Unlike MVA, the v-NY strain is a replication-competent virus derived from the New York City Board of Health viral strain of smallpox vaccine (44–47) that displays reduced virulence compared to the standard smallpox vaccine (Dryvax). Due to the different features of these two vaccinia virus strains, we tested different prime-boost combinations of both vaccines to establish an effective regimen for immunoactivation in C57BL/6 mice. Furthermore, we demonstrate that our vaccination regimens generated antibody and T cell responses in mice and protected Syrian hamsters from SARS-CoV-2 infection.

## Results

### Generation of recombinant v-NY-S and MVA-S viruses expressing full-length SARS-CoV-2 S protein

The recombinant vaccinia viruses MVA-S and v-NY-S were generated by inserting an ORF encoding full-length SARS-CoV-2 S protein (Wuhan-Hu-1, NC_045512) into the *tk* locus of the vaccinia virus strains MVA and v-NY, respectively (Fig. 1A). Cells infected with MVA-S and v-NY-S expressed high levels of SARS-CoV-2 S protein on cell surfaces, as revealed by flow cytometry (Fig. 1B) and immunofluorescence microscopy (Fig. 1C) analyses. Both full-length and processed forms of S protein were detected in immunoblots (Fig. 1D), confirming that the MVA-S and v-NY-S viruses stably expressed and processed SARS-CoV-2 S protein.

**Fig. 1.**
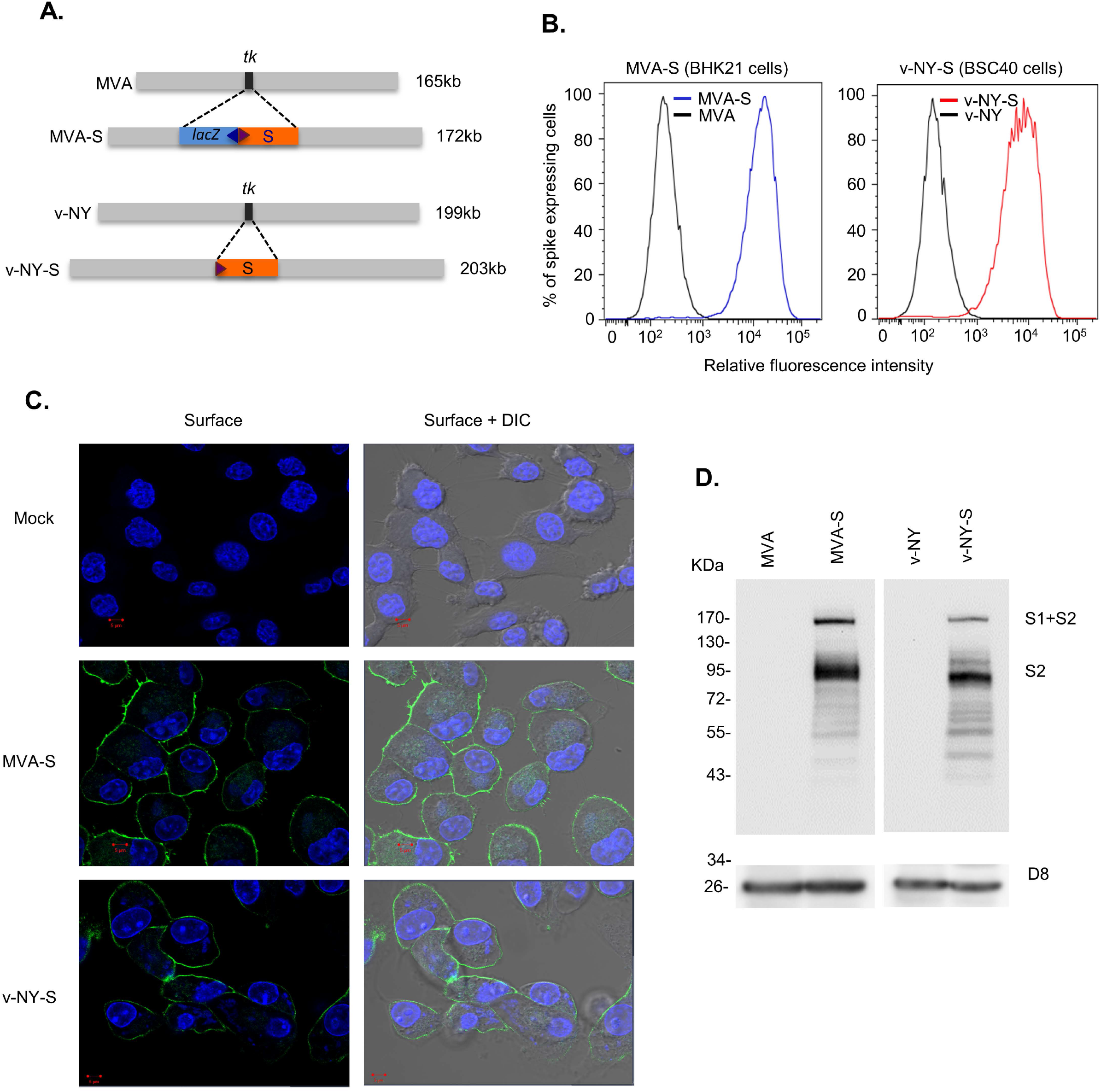
Generation and characterization of v-NY-S and MVA-S. **(A).** Schematic representation of the *tk* locus in the viral genomes of MVA-S and v-NY-S. The red box represents the ORF encoding SARS-CoV-2 S protein and the blue box represents the *lacZ* ORF. The small triangles represent viral promoters that drive gene transcription **(B).** Surface detection of SARS-CoV-2 S protein expressed from MVA- S and v-NY-S. BHK21 and BSC40 cells were infected with MVA-S (blue line) or v- NY-S (red line), respectively, harvested at 12 hours post-infection (hpi), stained with anti-RBD antibody, and then analyzed by flow cytometry. **(C).** Immunofluorescence staining of SARS-CoV-2 S protein in cells infected with MVA-S and v-NY-S. BHK21 and BSC40 cells were infected with MVA-S or v-NY-S at an MOI of 5 and fixed at 12 hpi with 4% paraformaldehyde, stained with anti-RBD antibody (green), and then photographed. Cell nuclei were stained with DAPI (blue). **(D).** Immunoblot of SARS-CoV-2 S protein expressed by MVA-S and v-NY-S. BHK21 cells were infected with MVA or MVA-S; BSC40 cells were infected with v-NY or v-NY-S, respectively, and harvested at 12 hpi for immunoblot analyses with anti-S2 antibody. Vaccinia D8 protein was used as a control.

### v-NY-S and MVA-S prime-boost vaccination regimens generate neutralizing antibodies in immunized C57BL/6 mice

We designed three prime-boost vaccination regimens using MVA-S and v-NY-S viruses (Fig. 2A). Control mice were primed and boosted with PBS buffer alone. For the first regimen, (MVA5/MVA1), mice were primed i.m. with 5×10^7^ PFU/animal of MVA-S virus and boosted with 1×10^7^ PFU/animal of MVA-S. For the second regimen, (vNY1/MVA1), mice were primed with 1×10^7^ PFU/animal of v-NY-S by means of tail scarification (t.s.) and then boosted i.m. with 1×10^7^ PFU/animal of MVA-S. For the third regimen, (vNY5/MVA1), mice were primed with v-NY-S at a higher titer of 5×10^7^ PFU/mouse by t.s. and then boosted i.m. with 1×10^7^ PFU/animal of MVA-S. For each regimen, the mice were primed at day 0 and primary (1°) sera were collected 4 weeks later. These mice were then rested for 3 days, boosted, and then secondary (2°) sera were drawn 2 weeks later. In some experiments, spleens were harvested 4 weeks after boosting for T cell and cytokine analyses. The t.s. site of vaccinated mice healed well and the mice remained healthy without any loss of body weight (S1Fig.). 1° and 2° sera were collected from mice and flow cytometry revealed that they recognized SARS-CoV-2 S protein expressed from recombinant S-BAC baculovirus (Fig. 2B). Quantification (Fig. 2C) confirmed that anti-spike antibodies were specifically generated after primary immunization and that antibody titers were significantly enhanced after vaccine boosting. Mice primed with v-NY-S presented higher levels of anti-spike antibody compared to those primed with MVA-S (Fig. 2C). Immunoblot analyses (Fig. 2D) also revealed anti-spike antibody reactivity to recombinant S protein, consistent with our FACS data (Fig. 2C). We tested the neutralizing activity of 2° sera using SARS-CoV-2 spike pseudotyped virus (Fig. 2E, panel i) and SARS-CoV-2 virus (Fig. 2E, panel ii). Neutralization activity is presented as the reciprocal dilution of serum required for 50% inhibition of virus infection (ND_50_). Our results show that all three regimens successfully generated high titers of neutralizing antibodies that inhibited SARS-CoV-2 S protein-mediated virus entry in both infection systems. Finally, we tested if our prime-boost vaccination regimens induced long-lasting antibody responses by comparing mouse sera collected at 0.5 and 4.5 months after the MVA5/MVA1 and vNY1/MVA1 regimens. Sera taken 4.5 months after boosting still contained 60-80% of spike-specific antibodies, as revealed by FACS analyses (Fig. 2F), and a pseudotyped SARS-CoV-2 virus infection assay demonstrated that they retained comparable neutralization activity to sera at 0.5 months (Fig. 2G), indicating these two vaccination regimens can elicit long-lived anti-spike antibody responses that have been shown to correlate with protection against SARS-CoV-2.

**Fig. 2.**
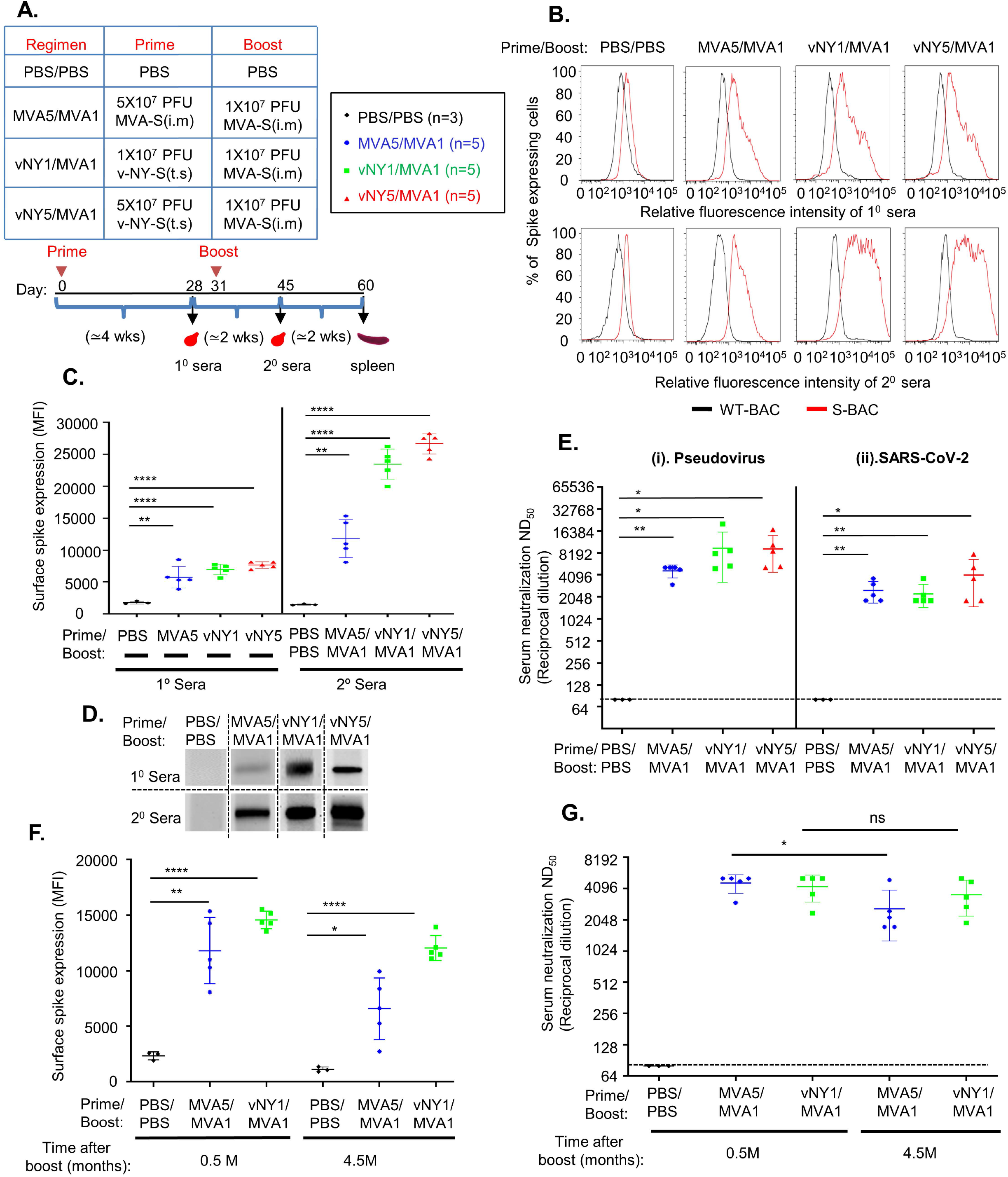
Prime-boost MVA5/MVA1, vNY1/MVA 1and vNY5/MVA1 vaccination regimens elicited SARS-CoV-2 S protein-specific neutralizing antibodies in C57BL/6 mice. **(A).** Summary and timeline of the three prime-boost vaccination regimens and analyses. **(B).** Primary and secondary sera from immunized mice recognized SARS-CoV-2 S protein on cell surfaces. Mouse sera collected 4 weeks after priming (1° sera) and 2 weeks after boosting (2° sera) were assessed for SARS- CoV-2-specific IgG antibodies by flow cytometry using SF9 cells infected with either S-BAC (red line) or WT-BAC (black line). A single representative serum is shown in each histogram. **(C).** Quantification of anti-spike antibody titers in 1° and 2° sera from mice (as shown in B) using the mean fluorescence intensity (MFI) value from FACS. Numbers of mice for 1° and 2° sera collection are identical: PBS vs. PBS/PBS control (n=3); MVA5 vs. MVA5/MVA1 (n=5); vNY1 vs. vNY1/MVA1 (n=5), and vNY5 vs. v-NY5/MVA1 (n=5). Data are represented as mean ± SD. **p<0.01; ***p<0.001; ****p<0.0001. **(D)**. Immunoblot analyses of recombinant SARS-CoV-2 S protein using 1° and 2° sera (1:100) from immunized mice. A single representative serum is shown in each blot. **(E).** Neutralization assays of 2° sera collected from vaccinated mice using (i) Pseudovirus and (ii) SARS-CoV-2 virus infection: PBS/PBS control (n=3); MVA5MVA1 (n=5); vNY1/MVA1 (n=5); and vNY5/MVA1 (n=5). The dotted line represents assay limits of detection. *p<0.05; **p<0.01. **(F).** Quantification of anti-spike antibodies in mouse sera collected at 0.5 and 4.5 months after vaccination regimens using SF9 cells infected with WT-BAC or S-BAC. *p<0.05; **p<0.01; ****p<0.0001. **(G)**. Pseudovirus neutralization assay using mouse sera collected at 0.5 months and 4.5 months after vaccination regimens: PBS/PBS control (n=3); MVA5/MVA1 (n=5); and vNY1/MVA1 (n=5). ns – not significant. *p<0.05. The dotted line represents assay limits of detection.

### v-NY-S and MVA-S immunization generates a T_H_1-biased immune response in mice

IFN-γ-producing T_H_1 cells promote a B-cell class switch towards IgG2a/IgG2c, whereas IL-4-producing T_H_2 cells promote a class switch towards IgG1 (49, 50). Therefore, a ratio of IgG2c (or IgG2a) to IgG1 >1 is a good indicator of a T_H_1-biased immune response, which is important for pathogen clearance. Accordingly, we used ELISA to measure levels of the IgG2c and IgG1 isotypes of anti-spike antibodies in C57BL/6 mouse sera collected after vaccination regimens (Fig. 3A). All three vaccination regimens induced production of the IgG2c and IgG1 isotypes (Fig. 3A) and with IgG2c/IgG1 ratios > 1 (Fig. 3B), suggesting they had elicited a T_H_1-biased immune response. We further *in vitro*-stimulated splenocytes from vaccinated mice with a SARS-CoV-2 spike peptide pool and then counted cells secreting T_H_1 cytokines (IL−2, IFN-γ and TNF-α) and T_H_2 cytokines (IL-4 and IL-6) (Fig. 3C) (51, 52). Consistently, we found that more cells secreted TNF-α and IFN-γ than IL-4 and IL-6 (Fig. 3C), supporting that our three vaccination regimens triggered a T_H_1-biased response(51, 52). Furthermore, we investigated if our immunization regimens generated T effector memory (Tem) cells that are known to play a critical role in immune protection against secondary viral infections in lung tissue (53). Splenocytes isolated from mice 4 weeks after vaccination regimens were incubated with a SARS-CoV-2 spike peptide pool for 2 h and then analyzed by flow cytometry (Fig. 3D & 3E), which revealed that all three regimens resulted in significantly increased numbers of CD8^+^ Tem cells (Fig. 3D), but not CD4^+^ Tem cells (Fig. 3E), in spleen tissue.

**Fig. 3.**
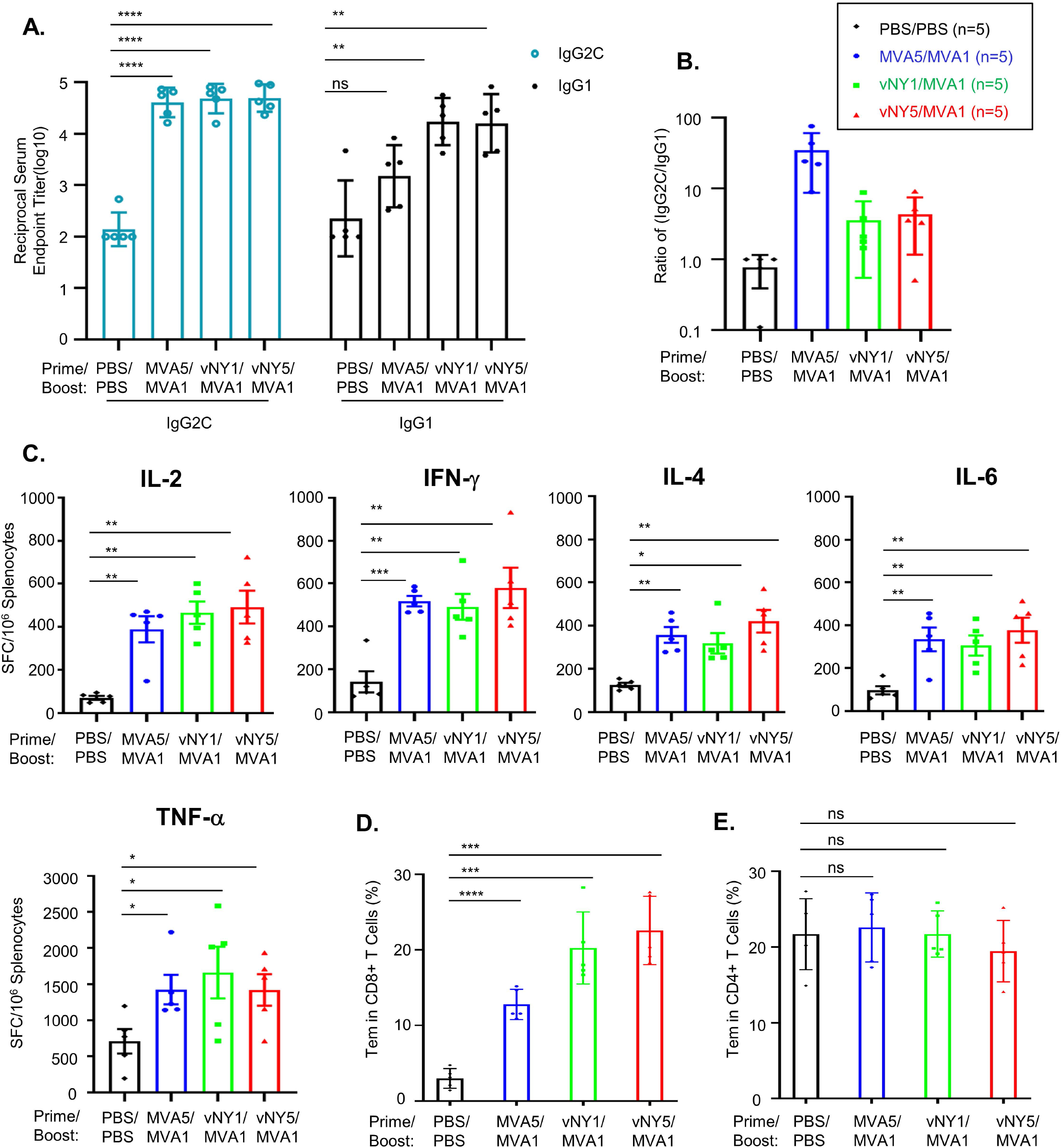
MVA5/MVA1, vNY1/MVA1 and vNY5/MVA1 vaccination regimens induce T_H_1-biased immune responses. **(A).** End-point titers of SARS-CoV-2 spike-specific IgG2C and IgG1 antibodies in mouse sera collected 2 weeks after vaccination regimens. ns-not significant; **p<0.01; ****p<0.0001. **(B).** End-point titer IgG2C/IgG1 ratio calculated based on data from (A) (n=5 for each group). **(C).** ELISpot analyses of mouse splenocytes collected 4 weeks after vaccination regimens for their expression of IL-2, IFN-γ, TNF-α, IL4 and IL6 cytokines (n=5 for each group). Data represented as mean ± SEM. SFC – spot-forming cells. *p<0.05; **p<0.01; ***p<0.001. **(D& E).** SARS-CoV-2 spike-specific CD8^+^ (in D) and CD4^+^ (in E) T effector memory cells (CD44^+^CD62L^-^) in splenocytes, as detected by flow cytometry (n=5 for each group). Data represented as mean ± SD. ns – not significant; ***p<0.001; ****p<0.0001.

### v-NY-S and MVA-S immunization generated neutralizing antibodies in immunized Syrian hamsters

C57BL/6 mice are not susceptible to SARS-CoV-2 infection, whereas Syrian hamsters serve as an appropriate animal model of respiratory infection by SARS-CoV-2 in human (54–65). We subjected Syrian hamsters to the same prime-boost vaccination regimens, i.e., MVA5/MVA1, vNY1/MVA1 and vNY5/MVA1 (Fig. 4A) as applied to mice (Fig. 2A), except that we used a skin scarification inoculation approach for hamsters. A small scar formed at the immunization site, which healed within two weeks (S2A Fig.), and the immunized hamsters remained healthy any did not exhibit weight loss (S2B Fig.). Primary and secondary sera collected from these immunized hamsters specifically recognized SARS-CoV-2 S protein expressed on cell surfaces (Fig. 4B). Quantification of all hamster sera by FACS demonstrated that boosting enhanced anti-spike antibody titers (Fig. 4C), which was confirmed by immunoblotting (Fig. 4D). Importantly, all three vaccination regimens generated anti-spike antibodies with high neutralization activity in both pseudotyped SARS-CoV-2 virus (Fig. 4E, panel i) and live SARS-CoV-2 virus infection assays (Fig. 4E, panel ii). Taken together, these data show that, as observed for mice, our vaccination regimens generated high antibody titers in hamsters that neutralized SARS-CoV-2 infection.

**Fig. 4.**
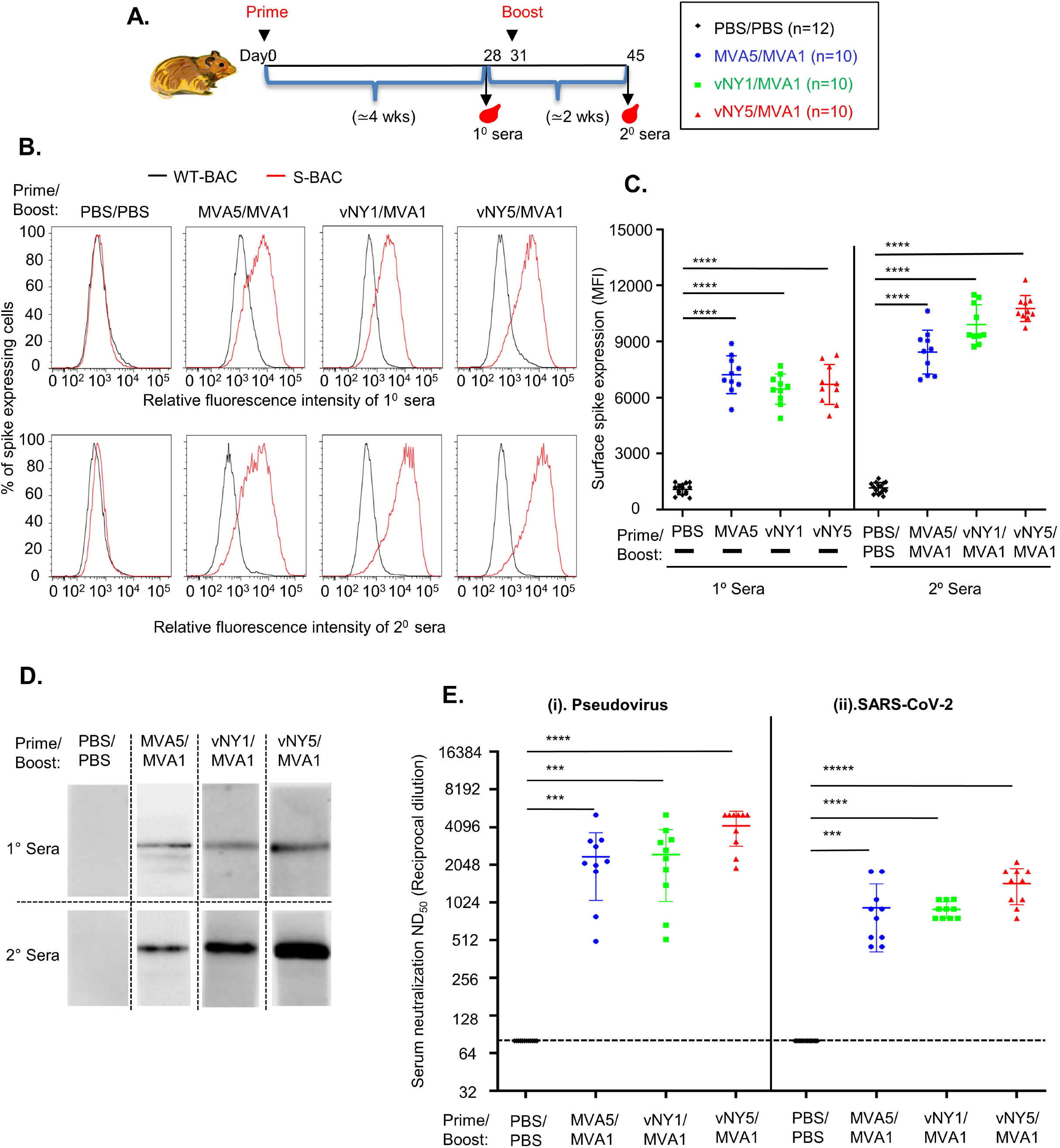
MVA5/MVA1, vNY1/MVA1 and vNY5/MVA1 prime-boost vaccination regimens generated SARS-CoV-2 spike-specific neutralizing antibodies in Syrian hamsters. **(A).** Timeline for hamster immunization and sera collection. **(B).** Primary and secondary sera from immunized hamsters recognized SARS-CoV-2 S protein on cell surfaces. Hamster sera collected 4 weeks after priming (1° sera) and 2 weeks after boosting (2° sera) were assessed for SARS-CoV-2-specific IgG antibodies by flow cytometry using SF9 cells infected with either S-BAC (red line) or WT-BAC (black line). A single representative serum is shown in each histogram. **(C).** Quantification of anti-spike antibody titers in 1° and 2° sera from hamsters in B, using the mean fluorescence intensity (MFI) value from FACS. Numbers of hamsters for 1° and 2° sera collection are identical: PBS vs. PBS/PBS control (n=15); MVA5 vs. MVA5/MVA1 (n=10); vNY1 vs. vNY1/MVA1 (n=10), and vNY5 vs. v-NY5/MVA1 (n=10). Data are represented as mean ± SD. ****p<0.0001. **(D).** Immunoblots of 1° and 2° sera (1:20) from immunized hamsters using recombinant SARS-CoV-2 S protein. A single representative serum is shown in each blot. **(E).** Neutralization assays of 2° sera collected from vaccinated hamsters using (i) pseudovirus and (ii) SARS-CoV-2 virus infection: PBS control (n=12); MVA5/MVA1 (n=10); vNY1/MVA1 (n=10); and vNY5/MVA1 (n=10). The dotted line represents assay limits of detection. ***p<0.001; ****p<0.0001.

### v-NY-S and MVA-S immunization reduce lung pathology in SARS-CoV-2- infected Syrian hamsters

Next, we performed challenge experiments in hamsters 2 weeks after vaccine boosting by intranasally (i.n) inoculating 1×10^5^ PFU/animal of SARS-CoV-2 virus into each hamster and then measuring changes in body weight till 3 d.p.i. (Fig. 5A). Control hamsters (immunized with a PBS placebo) presented minor but detectable weight loss at 3 d.p.i., whereas those subjected to our vaccination regimens presented no obvious weight loss (Fig. 5B). Previous studies showed that SARS-CoV-2 infection of Syrian hamsters results in virus replication in lung tissue and that virus titers often peaked from 2-4 d.p.i. and gradually cleared by 7 d.p.i. (54, 56, 58, 66–70). Therefore, we sacrificed hamsters at 3 d.p.i. and then measured SARS-CoV-2 virus titers in their lungs (Fig. 5C). None of the MVA5/MVA1- or vNY1/MVA1-vaccinated hamsters presented detectable levels of SARS-CoV-2 virus in their lung tissue, whereas virus titers of up to ∼ 4×10^6^ TCID_50_ were detected in the lungs of the placebo group (Figure 5C). Moreover, no virus was detected in nine vNY5/MVA1-immunized hamsters and only one such animal presented residual amounts of virus (< 0.1% of the mean virus titer of the placebo group) (Fig. 5C).

**Fig. 5.**
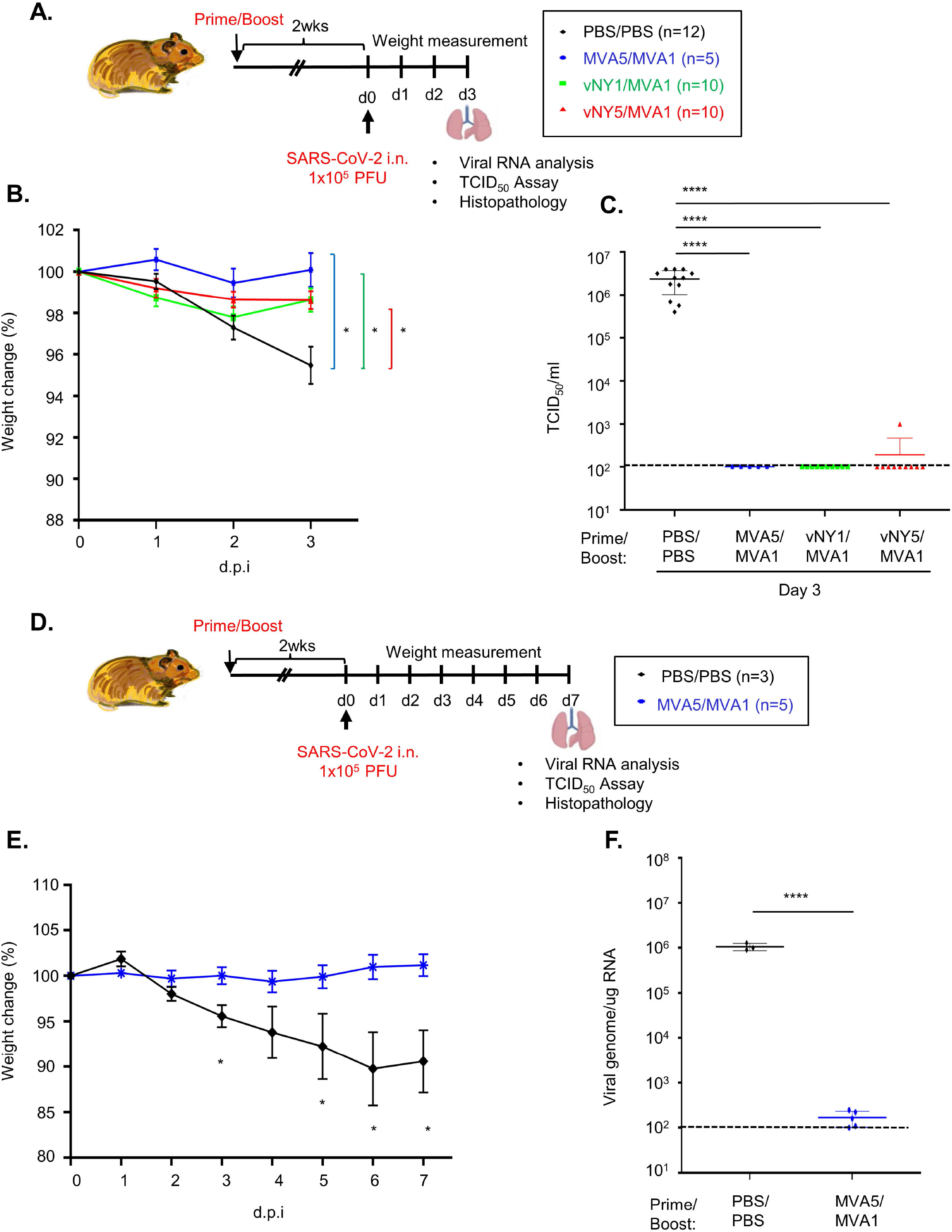
Hamsters subjected to the MVA5/MVA1, vNY1/MVA1 or v-NY5/MVA1 vaccination regimens were protected against intranasally-administered SARS- CoV-2 infection. **(A).** Timeline of the immunization and challenge experiments. Hamsters immunized with one of three prime-boost vaccination regimens (MVA5/MVA1, vNY1/MVA1 or vNY5/MVA1), or placebo (PBS) as a control, were challenged i.n. with 1×10^5^ PFU SARS-CoV-2 virus, before harvesting lungs at 3d.p.i.. **(B).** Weight change of hamsters within 3 days of SARS-CoV-2 challenge. Data represented as mean ± SEM. *p<0.05. **(C).** TCID_50_ value of SARS-CoV-2 in lung tissue of hamsters at 3 d.p.i. after SARS-CoV-2 challenge: PBS/PBS control (n=12); MVA5/MVA1 (n=5); vNY1/MVA1 (n=10); and vNY5/MVA1 (n=10). Data represented as mean ± SD. ****p<0.0001. **(D).** Timeline of the immunization and challenge experiments. Hamsters immunized with one prime-boost vaccination regimen (MVA5/MVA1), or placebo (PBS/PBS) as a control, were challenged i.n. with 1×10^5^ PFU SARS-CoV-2 virus, before harvesting lungs at 7d.p.i.. **(E)** Weight change of hamsters within 7 days of SARS-CoV-2 challenge: PBS/PBS control (n=3); and MVA5/MVA1 (n=5). Data represented as mean ± SEM. *p<0.05. **(F).** SARS-CoV-2 genomic RNA in lungs of MVA5/MVA1-immunized hamsters at 7 d.p.i. after SARS-CoV-2 challenge: PBS/PBS control (n=3); and MVA5/MVA1 (n=5). Unless stated otherwise, data are represented as mean ± SD. The dotted line represents assay limits of detection. ****p<0.0001.

We further explored the impact of our MVA5/MVA1 regimen at 7 d.p.i. (Fig.5D). The weight loss of the placebo group was even more pronounced at 7 d.p.i. than at 3 d.p.i. (∼10-15%), whereas that of MVA5/MVA1- immunized hamsters remained unchanged (Fig. 5E). When we harvested lungs from immunized or placebo hamsters at 7 d.p.i. and measured SARS-CoV-2 virus titers and viral RNA levels, we found that no virus was detected in any of the hamsters (S3 Fig.), but ∼10^6^ copies of viral RNA were detected in the lungs of the placebo group, whereas only ∼10^2^ copies were detected in MVA5/MVA1-immunized hamsters (Fig. 5F).

To further validate our findings, we removed the lungs of experimental hamsters at 3 and 7 d.p.i. and processed them for histological examination (Fig. 6). The lungs of placebo-infected hamsters at 3 d.p.i. presented diffuse congestion, shrinking of alveoli, hemorrhaging, and mononuclear cell infiltration (Fig. 6A, open arrowheads). Moreover, bronchiolar epithelia vacuolization, necrosis and inflammatory exudates were also observed, and there was pronounced vasculitis and/or endothelialitis (Fig. 6A, black arrows) involving both medium and small blood vessels disrupted by a mixture of immune infiltrates. Immunostaining with an antibody against SARS-CoV- 2 nucleocapsid (NP) protein revealed some areas of peribronchiolar immunoreactivity (Fig. 6A), mainly in the pneumocytes and less commonly in bronchiolar epithelial cells. Interestingly, despite the severe endothelial destruction observed by H&E staining, the SARS-CoV-2 NP antibody we deployed did not detect any positive viral-protein signal in the blood vessels of these placebo-infected hamsters. In contrast to the striking bronchointerstitial pneumonia observed in placebo-infected hamsters, there was only minimal to mild lung inflammation at 3 d.p.i. in the hamster groups subjected to the three vaccination regimens and SARS-CoV-2 NP protein signal was barely detectable (Fig. 6B-D).

**Fig. 6.**
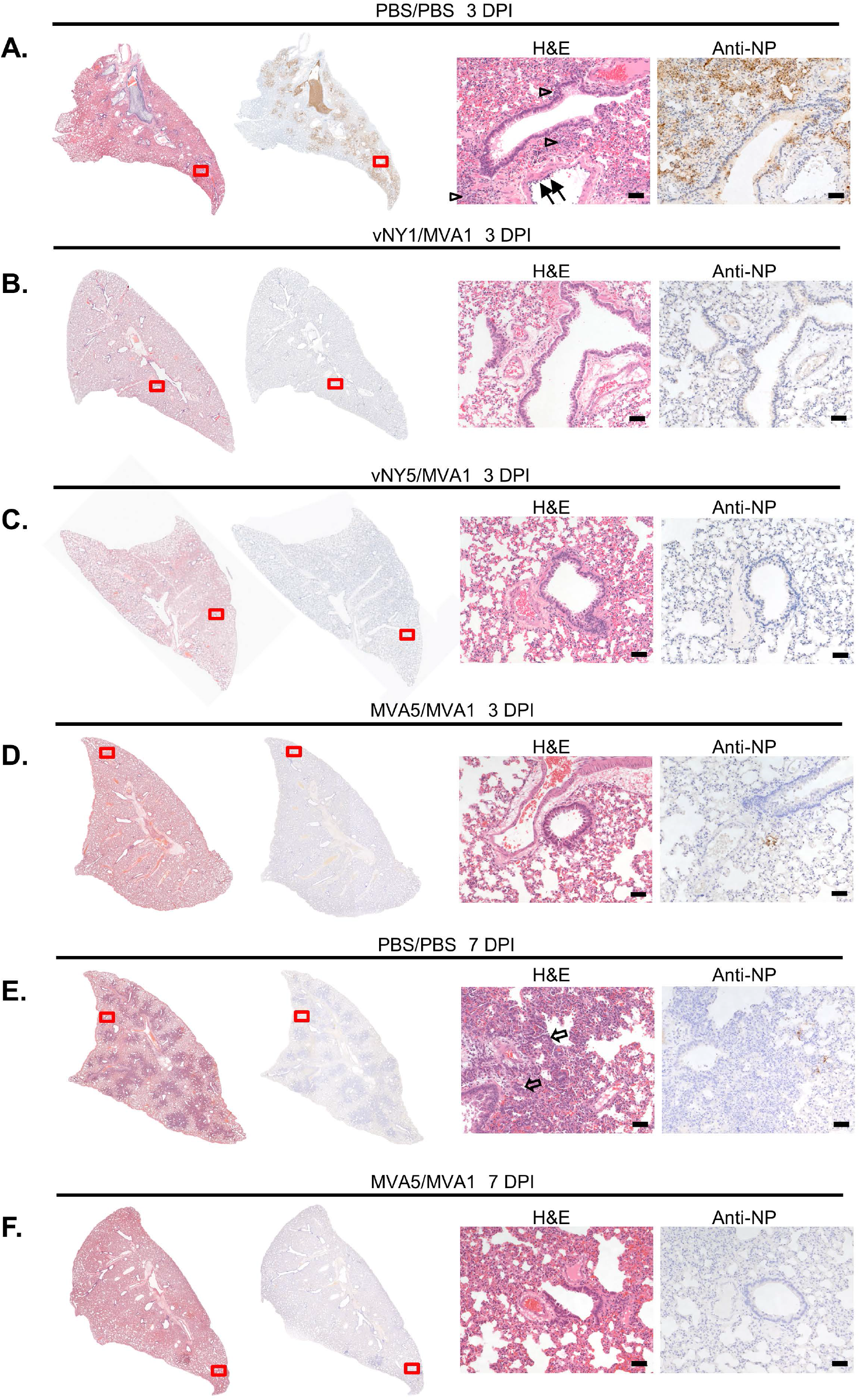
Lung pathology and immunohistochemistry of hamsters after SARS-CoV-2 challenge. **(A).** H&E and immunohistochemical staining of lungs of the placebo (PBS/PBS) infection hamster group at 3 d.p.i.. H&E staining showed severe bronchointerstitial pneumonia with the alveolar walls thickened by edema, capillary congestion and variable immune cell infiltration (open arrowheads). The vascular endothelia were frequently disrupted by immune infiltrates (vasculitis/endotheliolitis, black arrows). Immunohistochemistry of SARS-CoV-2 NP protein revealed prominent peribronchiolar staining (arrow heads). **(B,C & D).** H&E and immunohistochemical staining of lungs from the vNY1/MVA1 (B), vNY5/MVA1 (C) and MVA5/MVA1 (D) hamster groups at 3 d.p.i.. Compared to the placebo (PBS/PBS) infection hamster group, lung architecture was better preserved, there was much less immune cell infiltration, and SARS-CoV-2 NP staining signal was barely detectable. **(E).** H&E and immunohistochemical staining of lungs of the placebo (PBS/PBS) infection hamster group at 7 d.p.i.. H&E staining revealed prominent type II pneumocytic hyperplasia (open arrows) with variable immune cell infiltration. Immunohistochemistry of SARS-CoV-2 NP protein detected dispersed positive signals at the edges of regenerative foci. **(F).** MVA5MVA1-immunized hamsters displayed minimal lung pathology and scant SARS-CoV NP immunolabeling at 7 d.p.i.. The enlarged views of H&E and immunohistochemistry-stained regions are marked by red boxes. The scale bar represents 50 μm.

We also examined the lung tissues of hamsters of the placebo and MVA5/MVA1 groups at 7 d.p.i. (Fig. 6E & 6F). Profound type II pneumocyte hyperplasia (Fig. 6E, open arrows) was observed for the placebo-infection group, accompanied by mild to moderate neutrophilic infiltrate and numerous megakaryocytes centered on an obliterated bronchiole. Immunohistochemistry revealed weak but positive anti-NP antibody signal in pneumocytes at the periphery of bronchiole-centered lesions of placebo-infected hamsters (Fig. 6E). In contrast, the lungs of the MVA5/MVA1- infected group presented a less inflammatory phenotype at 7 d.p.i. and barely detectable anti-NP signal. Thus, taken together, our prime-boost vaccination regimens prevent SARS-CoV-2 viral spread in lung tissues and reduce inflammation and lung pathology.

### Single immunization with v-NY-S partially protects Syrian hamsters from SARS-CoV-2 infection

We wished to establish if single-dose immunization with recombinant v-NY-S virus could provide protection against SARS-CoV-2 in Syrian hamsters. Hamsters were immunized with PBS (placebo), 1×10^7^ or 5×10^7^ PFU/animal of v-NY-S by skin scarification and then sera were collected 2 weeks later (Fig. 7A). The sera were subjected to a SARS-CoV-2 pseudovirus neutralization assay, which showed that priming with v-NY-S alone generated neutralizing antibodies against SARS-CoV-2 virus in a dosage-dependent manner (Fig. 7B). Then we performed challenge experiments and monitored SARS-CoV-2 virus titers in lungs at 3 d.p.i. (Fig. 7C). Virus titers were ∼10^6^ PFU/animal in the placebo-infected group, but virus titers were >100-fold lower in hamsters subjected to single immunization with v-NY-S at either dosage (1×10^7^ or 5×10^7^ PFU/animal), showing that single immunization had already provided partial protection against SARS-CoV-2 infection. Upon removing lungs for histological examination, we observed that the placebo-infected group presented a severe pathological phenotype including diffuse congestion, shrinking of alveoli, hemorrhaging, and mononuclear cell infiltration (Fig. 7D, open arrowheads).

**Fig. 7.**
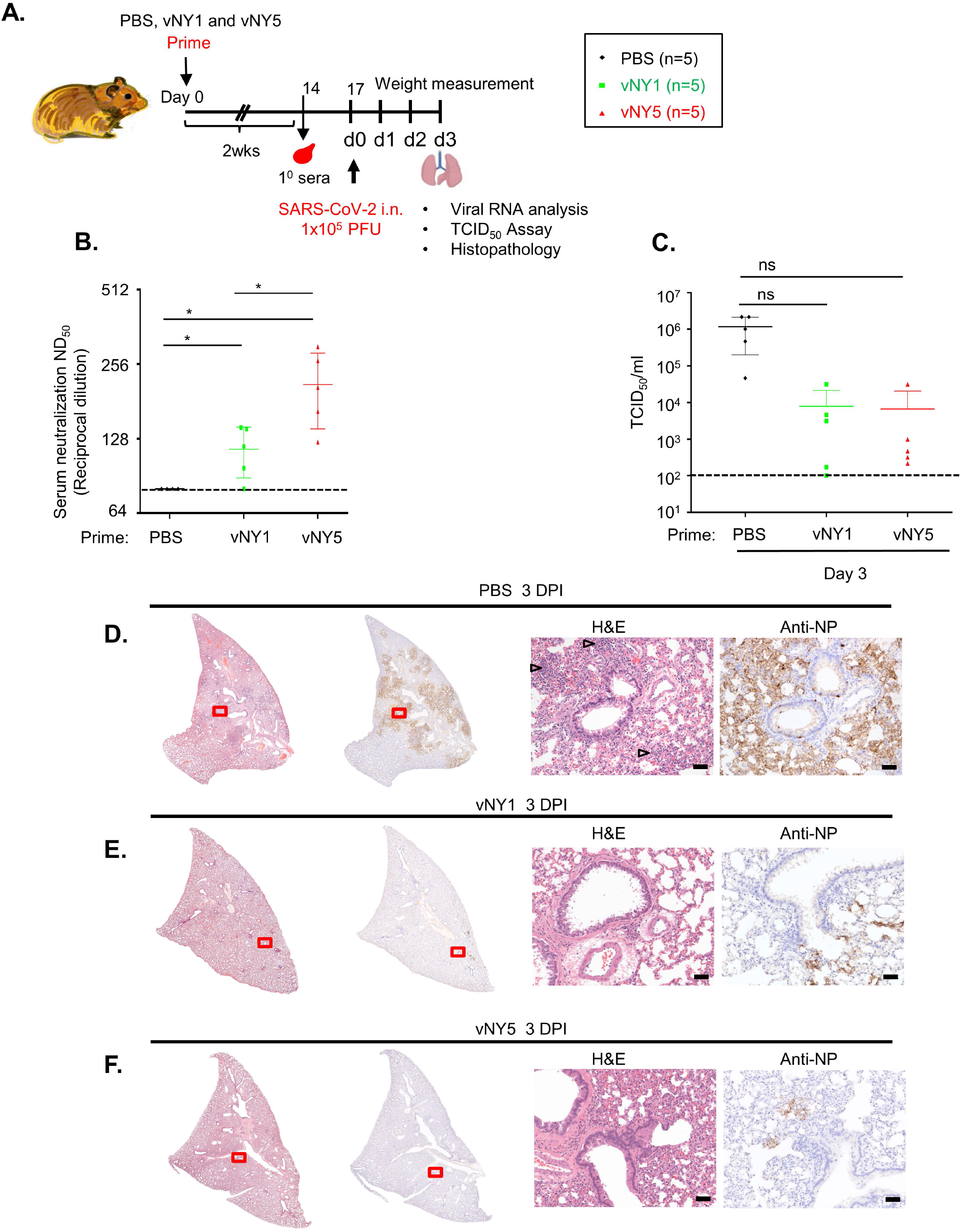
Single-dose vNY1 or vNY5 vaccination partially protected hamsters from intranasally-administered SARS-CoV-2 infection. **(A).** Timeline showing the immunization and challenge experiment. Hamsters immunized with a single dose of vNY1, vNY5 or placebo (PBS) were challenged i.n. with 1×10^5^ PFU SARS-CoV-2 virus and then lungs were harvested at 3 d.p.i.. **(B).** Pseudovirus neutralization assays of the 1° sera collected 2 weeks after vaccine priming in hamsters. PBS control (n=4); vNY1 (n=5); and vNY5 (n=5). Data represented as mean ± SD. The dotted line represents assay limits of detection. *p<0.05. **(C).** TCID_50_ values for lungs from hamsters of the placebo (PBS), vNY1 and vNY5 groups at 3 d.p.i. after SARS-CoV-2 challenge (n=5 for each group). Data represented as mean ± SD. The dotted line represents assay limits of detection. ns- not significant. **(D).** H&E and immunohistochemical staining of lungs of the placebo (PBS) infection hamster group at 3 d.p.i.. H&E staining revealed an identical pathology to that shown in Figure 6A, revealing severe bronchointerstitial pneumonia with the alveolar walls thickened by edema, capillary congestion and variable immune cell infiltration (open arrowheads). Immunohistochemistry of SARS-CoV-2 NP protein revealed prominent peribronchiolar staining (arrowheads), with the vascular endothelia frequently being disrupted by immune infiltrates. **(E and F).** H&E and immunohistochemical staining of hamster lungs primed with vNY1 (in E) or vNY5 (in F) at 3 d.p.i.. The lung architecture was largely preserved, displaying reduced immune cell infiltration relative to the placebo infection group and SARS-CoV-2 NP protein was barely detectable by immunohistochemistry. The scale bar represents 50 μm.

Moreover, immunostaining for SARS-CoV-2 NP protein also revealed widespread peribronchiolar immunoreactivity (Fig. 7D, arrowheads) in the lungs of the placebo group. In contrast, the lung pathology of the vNY1-infected (Fig. 7E) and vNY5- infected (Fig. 7F) groups was much milder than observed for the placebo-infected group, displaying lower immune cell infiltration, rare epithelial degeneration and an absence of vasculitis/endothelialitis. Viral NP immune signal in lung tissues was also significantly lower in the vNY1 and vNY5 groups relative to the placebo group (rightmost panels in Fig. 7E & 7F).

### Sera from vaccinated mice partially cross-neutralize SARS-CoV-2 variants of concern

Recently, circulation of new SARS-CoV-2 variants of concern (VOC) has become prevalent, entailing a risk of increasing resistance to the neutralizing antibodies generated by SARS-CoV-2 vaccines. Accordingly, we used pseudotyped SARS-CoV- 2 virus neutralization assays to test sera from our prime-boost vaccinated mice for their ability to cross-neutralize SARS-CoV-2 variants (Fig. 8). Although statistical analyses did not reveal a significant neutralization difference among the three immunization regimens to each variant, the mean neutralization titers of the α and γ variants were similar to that of the wild type whereas the mean titers of β, B.1.617 and δ variants appeared much lower than the wild type (Fig. 8A). For sera generated according to each regimen (Fig. 8B-D), we divided the value of ND_50_ for wild type by that of each variant to obtain the fold decrease in neutralization for each variant relative to wild type (Fig. 8B-D). We noticed that vNY5/MVA1 regimen maintained the least decrease of neutralization towards all variants (Fig. 8D) followed by vNY1/MVA1(Fig. 8C) and MVA5/MVA1 (Fig. 8B). Overall, antibodies from mice vaccinated with the heterologous prime-boost regimens, vNY1/MVA1 and vNY5/MVA1, appeared to display better cross-neutralization against different variants than those with the homologous prime-boost regimen (MVA5/MVA1).

**Fig. 8:**
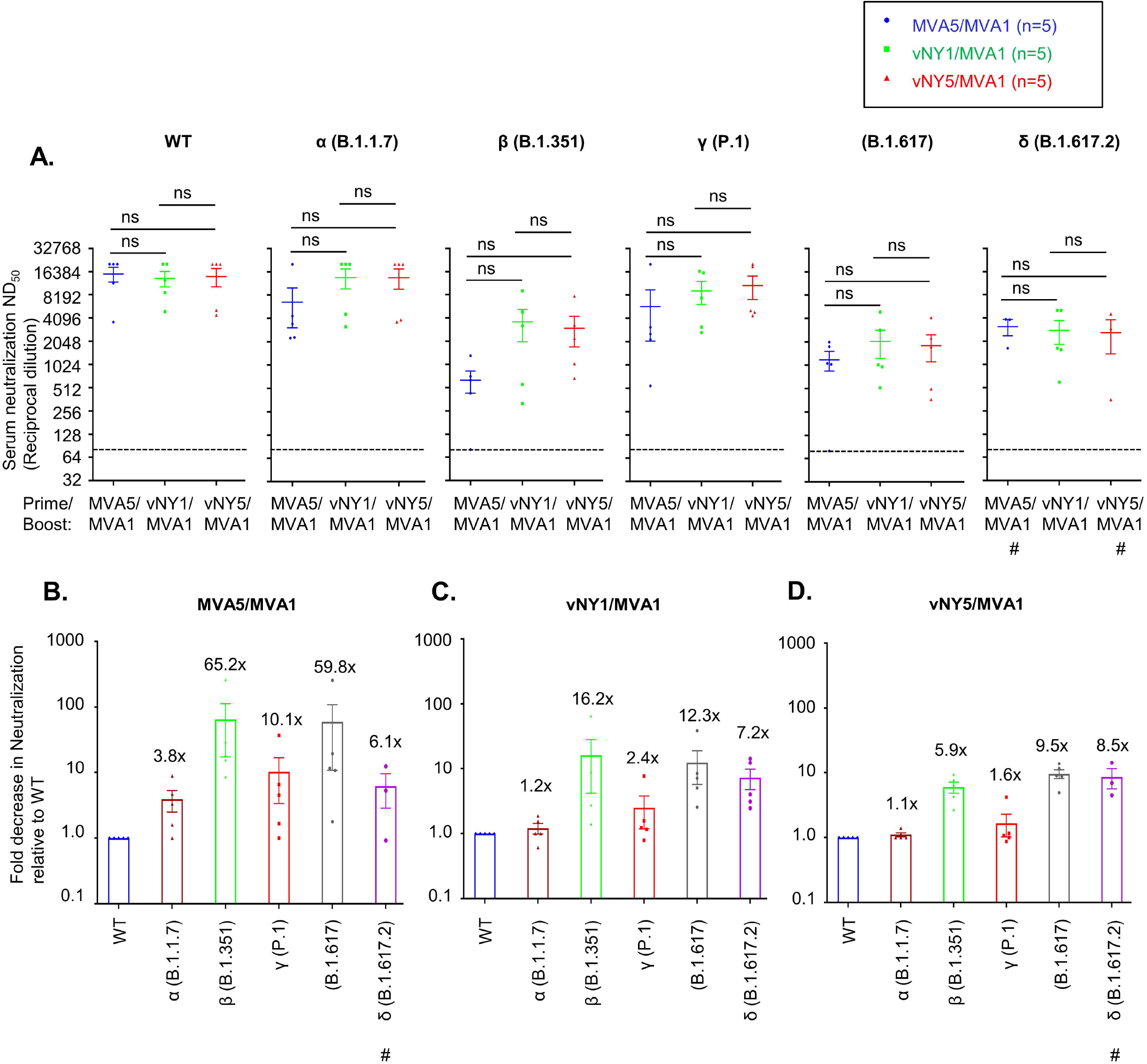
Sera from vaccinated mice partially cross-neutralized SARS-CoV-2 variants of concern. **(A).** Neutralization assays of secondary sera collected from vaccinated mice (regimens MVA5/MVA1, vNY1/MVA1 and vNY5/MVA1) following SARS-CoV-2 pseudotyped virus with spike protein from WT and variants α (B.1.17), β (B.1.351), γ (P.1), B1.617 and δ (B.1.617.2). The dotted line represents assay limit of detection. # represents n=3. ns-not significant. **(B-D).** Fold decrease for immunized sera from MVA5/MVA1 (B), vNY1/MVA1(C) and vNY5/MVA1(D) vaccination regimens in neutralization assays of individual variants relative to WT pseudovirus. Fold change was calculated by dividing the ND_50_ titer of each immunized serum against WT pseudovirus with the ND_50_ titer of the same serum against variant pseudovirus. # represents n=3. The dotted line represents assay limit of detection. Data are represented as mean ± SEM.

### A heterologous prime-boost immunization regimen with v-NY-S generates higher titers of neutralizing antibodies than a homologous prime-boost regimen

Our current regimens comprise both homologous (MVA5/MVA1) and heterologous (vNY1/MVA1 and vNY5/MVA1) prime-boost combinations, with these latter generating strong and long-lasting neutralizing antibody responses (Figure 2F). We wanted to explore other prime-boost combinations involving v-NY-S as an alternative strategy to improve immune responses. As shown in Fig. 9A, unlike MVA5/MVA1 (Figure 2C), prime-boosting with a homologous vNY1/vNY1 strategy did not increase anti-spike antibody titers after boosting (Fig. 9A). We rationalize that homologous prime-boosting with a replicating virus such as v-NY-S is not a good immunization strategy. Interestingly, heterologous priming with v-NY-S at 1×10^7^ PFU, followed by a boost with 5 μg recombinant spike protein (rS) plus adjuvant, enhanced anti-spike antibody titers (Figure 9A) and neutralizing activity (Figure 9B). In addition, we compared the route of priming between skin scarification (vNY1/MVA1) and intranasal inoculation (vNY0.1/MVA1) for v-NY-S and found that the latter route generated higher anti-spike antibody levels (Figure 9A) and neutralizing activity (Figure 9B) with only 1/10 of virus inoculum. These data are very encouraging and will aid in future vaccination strategies. Discussions are ongoing as to whether heterologous prime-boosting of COVID vaccines elicits more potent immune responses than a homologous prime-boost strategy (71–73). We argue that mix-and-match vaccination strategies are an important consideration for designing the next generation of SARS-CoV-2 vaccines.

**Fig. 9:**
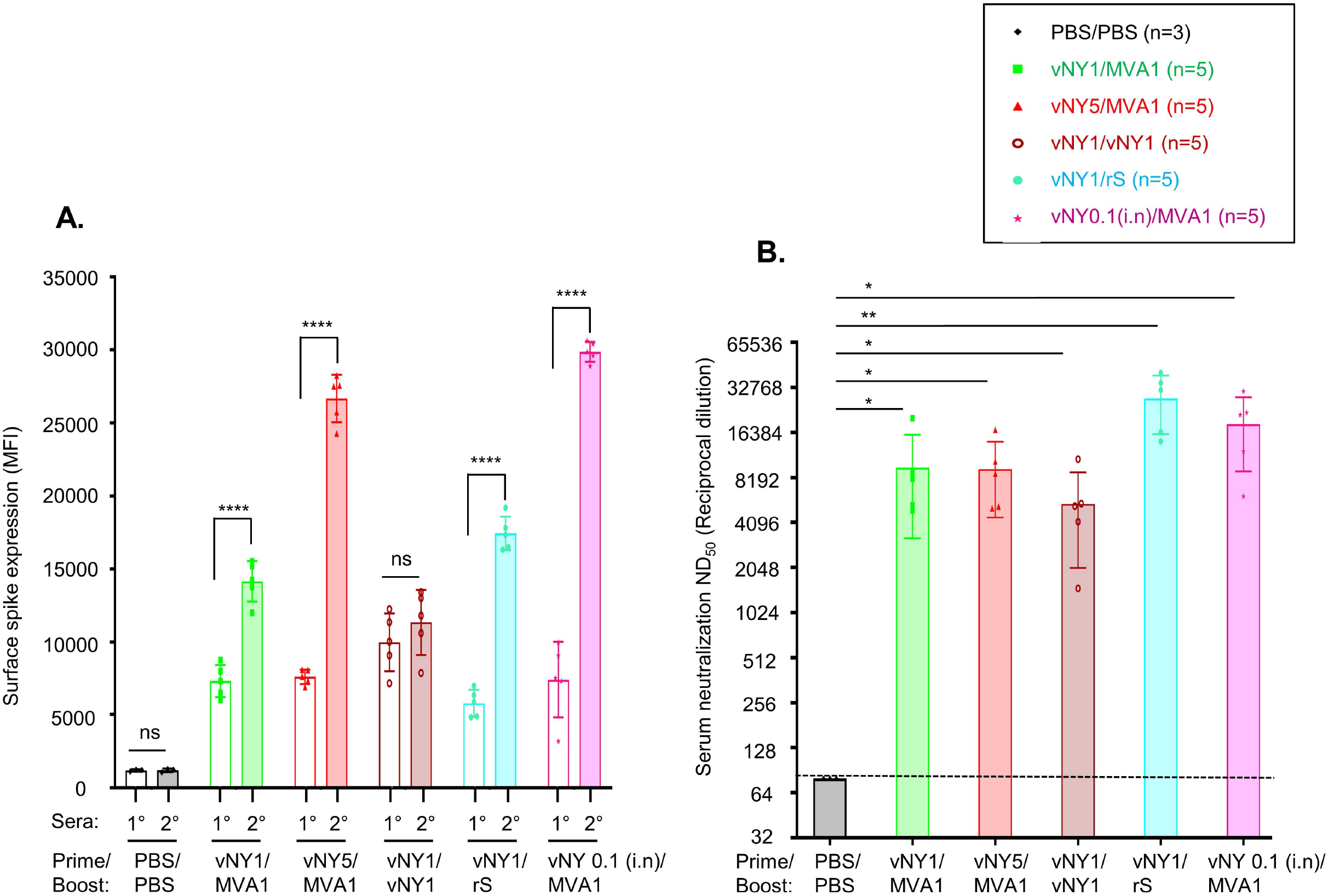
Heterologous prime-boost and intranasal inoculation with vNY-S enhances production of SARS-CoV-2 spike-specific neutralizing antibodies. **(A).** Quantification of anti-spike antibody titers in primary (1°, transparent bars) and secondary (2°, filled bars) sera from mice by staining cell surface spike protein on insect cells using FACS, as shown by the mean fluorescence intensity (MFI). Sera were collected from identical mice after prime (1°) and boost (2°). Numbers of mice in each prime-boost groups are shown on top of the figure. Data are represented as mean ± SD. ****p<0.0001; ns-not significant. **(B).** SARS-CoV-2 pseudotyped virus neutralization assays for 2° sera collected from mice after different prime-boost regimens. The dotted line represents assay limit of detection. rS: recombinant spike protein. *p<0.05; **p<0.01.

## Discussion

In this study, we have demonstrated in preclinical models the safety and immunogenicity of recombinant vaccinia viruses expressing SARS-CoV-2 virus spike protein. All three of the prime-boost vaccination regimens with v-NY-S and MVA-S recombinant viruses elicited strong and long-lasting neutralizing antibody responses against SARS-CoV-2, and also generated a T_H_1-biased T cell response that can promote pathogen clearance. Importantly, we have demonstrated that our vaccination regimens protected Syrian hamsters (representing an appropriate animal model of SARS-CoV-2-induced respiratory disease in human) from weight loss, elicited rapid clearance of SARS-CoV-2 virus, and reduced immune infiltrates in lung tissue.

Previous studies have revealed that virus shedding is often associated with a high viral titer in lung tissues (42, 54, 68). Bricker et al. (2021) showed that virus titers in vaccinated hamsters were cleared by day 3, and no virus shedding was detected in nasal swabs even as early as day 2 (68). Furthermore, Liu et al. (2021) found that vaccinated transgenic mice did not shed virus as early as day 2 when virus was cleared from the lungs (42). Since virus was cleared from the lung tissue of our prime-boost vaccinated animals, it is likely that virus shedding in these animals is limited.

When we compared data from male versus female hamsters in terms of weight loss and virus replication in lungs, we did not observe noticeable sex-associated differences in these hamsters after challenge with SARS-CoV-2 virus. Interestingly, Dhakal et al. (2021) reported that hamsters of both sexes exhibited similar weight loss, virus titers in lungs and cytokine response in the first 5-7 days after SAS-CoV-2 challenge (74). However, they also showed that, starting from day 7 up to 4 weeks, female hamsters developed greater antibody responses and recovered from weight loss faster than male hamsters (74). Because all our experiments were terminated within 1 week after SARS-CoV-2 challenge it could explain a lack of sex-associated difference in our experiments.

Hamster is a good animal model of SARS-CoV-2 with several advantages. It is suitable to study virus replication and transmission by contact and aerosols (75, 76). Most of hamsters infected with SARS-CoV-2 do not die, which is consistent with human infections, and the pathological features of the lungs in the infected animals resemble those observed in COVID-19 patients (77–79). Hamster is relatively low cost when compared with other animal models, such as ferrets (80) and non-human primates (81), and has been shown to mount an effective antibody response that clears virus within one week, making it a suitable model to study vaccine efficacy and antiviral drugs (56, 75, 82). On the other hand, several limitations of the hamster model exist. SARS-CoV-2 infection in humans affects multiple organ systems including the heart, kidneys, liver, spleen and large intestine (83, 84) whereas in hamsters live virus mainly replicates in respiratory tracts with little evidence of infectious virus in other organs, despite the detection of viral RNA in non-respiratory tissues (55, 85). Furthermore, a lack of hamster-specific biological reagents makes it difficult to characterize immune responses and the mechanisms of pathogenesis in hamsters.

A successful vaccine against SARS-CoV-2 should stimulate both humoral and cellular immune responses to inhibit virus replication and prevent disease in the host (86). In both mouse and hamster model our prime-boost immunization with MVA-S and v-NY-S generated high titers of neutralizing antibodies. Evidence from preclinical studies in nonhuman primates and hamsters indicted that vaccine-induced SARS-CoV-2 neutralizing antibodies correlate with protection against lung infection and clinical disease (19, 55, 87). While it is difficult to study T-cell immune response in hamsters, our immunization regiments induced T_H_1-biased immune response and effector memory CD8^+^T cells in mice, consistent with other studies using MVA-based SARS-CoV-2 vaccines (41–43, 88). Interestingly, Tan, et al (2021) reported that early induction of interferon-γ secreting SARS-CoV-2 specific T cells correlates with mild disease and rapid virus clearance, implying an important role of T cell immunity in host protection (89). How to improve a SARS-CoV-2 vaccine to induce a coordinated B and T responses will be of great importance in the future.

SARS-CoV-2 vaccines developed by Moderna, Pfizer/BioNTech, Novavax and Janssen Pharmaceutical companies expressed a designed SARS-CoV-2 spike protein, S-2P, which contain two proline substitutions at K986 and V987 to stabilize the prefusion conformation of spike protein (19, 87, 90, 91). It is thought the S-2P design could improve protein stability and elicit high titers of neutralizing antibodies (19, 87, 90, 91). We did not modify the spike protein in our recombinant virus constructs primarily because of two reasons: (i) Vaccinia viral promoters are known to be much stronger than cellular promoters (92, 93) so the spike protein abundance and stability may not be a concern. (ii). We think that the natural spike protein conformation may provide a large repertoire of antigenic epitopes to generate neutralizing antibodies against SARS-CoV-2 virus. Our neutralizing antibodies induced strong neutralizing titers, supporting that rationale. Besides, Tscherne eta al (2021) constructed a vaccinia MVA-SARS-2-S vaccine that expresses a native SARS-CoV-2 spike protein and their results also showed high neutralizing antibodies were induced (88). Moreover, the AstraZeneca vaccine, which lacks pre-fusion 2P stabilization, provided sufficient protection during clinical trials (94, 95). It is intriguing that a new version of spike modification known as HexaPro, that contains six proline substitutions to further stabilize the pre-fusion S protein, was reported (96). Whether HexaPro represents a promising antigen for vaccine design will be interesting to know in the future.

While we were testing our vaccination regimens, several research groups published studies showing that MVA-based vaccines provided protection against SARS-CoV-2 infection in hACE2 transgenic mice (41, 88, 97) and macaques (43). The results of our recombinant MVA-S-based vaccines are consistent with those findings. Our study further explores the utility of the replication-competent v-NY strain, revealing that it could prove just as promising as the MVA strain in tackling SARS-CoV-2. The MVA strain that is growth-restricted in mammalian cells has been widely used in vaccine clinical studies due to its safety features. In contrast, v-NY was originally isolated and developed as a replicating vaccinia vector (98). A plaque-purified New York City Board of Health strain of vaccinia virus was established from a commercial preparation of smallpox vaccine (Dryvax) via three successive rounds of plaque- purification in BSC-40 cells. A master seed stock of this virus was prepared in human diploid MRC-5 cells, which was then used as the parental virus (v-NY) to construct a recombinant virus expressing the envelope glycoproteins of HIV-1 (HIVAC-1e) later used for FDA-approved clinical trials (44). Since then, master seed stocks of v-NY have been extensively characterized in terms of sterility, mycoplasma, adventitious viruses, plaque morphology, neutralization by vaccinia-specific monoclonal and polyclonal antisera, neurovirulence and genome restriction analysis (44). Unlike NYVAC (99), which is a highly attenuated virus constructed from the Copenhagen strain of vaccinia virus by specific genetic deletions, v-NY is a replication-competent virus and has not gone through any “attenuation protocol”, such as the repeated passages for MVA. The *in vitro* and *in vivo* characteristics of v-NY are similar to its parent virus, the New York City Board of Health strain of smallpox vaccine (44–47). In terms of safety profile, the recombinant v-NY virus (HIVAC-1e) has been tested in Phase I and II studies (46, 47). Moreover, v-NY has been used as a control virus in many animal studies and, in all cases, the elicited responses appear very similar to how smallpox vaccines behave in humans (44, 45, 100, 101).

There are several reasons why we elected to conduct v-NY-S inoculation via skin scarification. First, Langerin^+^ dermal dendritic cells have been shown to effectively present antigen upon skin scarification with vaccinia virus (102). Secondly, more antibodies are generated upon scarification than via intramuscular inoculation (103). Thirdly, given experience with the global smallpox vaccination program, skin scarification by means of a bifurcated needle to prick the skin 15 times within a few seconds is a convenient and efficient way to inoculate a large human population during a pandemic. It is possible that for people displaying acute skin breaks such as acne or suffering chronic skin conditions such as eczema or atopic dermatitis, the side-effects may be more serious and should be taken into consideration. In addition, the inoculation site on the arm should be properly cared for so that the virus does not spread. Overall, the availability of both recombinant MVA and v-NY viruses expressing SARS-CoV-2 S protein allows studies to determine whether genetic properties of the viral vector may modulate immune responses leading to differential vaccine efficacies. To this end, it may be worth thinking that a better ‘placebo vaccine’ control than PBS alone would be supernatants of cells that do not express the spike construct but that are prepared the same way as the vaccine. These preparations would contain a secretome including extracellular vesicles and other xenogeneic materials that might contribute to responses like cytokines, albeit presumably non-specific. It will be useful to include such controls in the future experiments.

Most SARS-CoV-2 vaccines currently on the market require a two-dose prime-boost vaccination program to generate sufficient protective immunity(12, 18, 20, 23, 104–109). Several Adenovirus-based vaccines (such as ChAdOx) were demonstrated as quite effective following two dosages (19, 20, 55, 110–112), though single immunization is sufficient for the Ad26.COV2.S vaccine (19). Because the fast rise of SARS-CoV-2 variants whether these vaccines can adequately control the COVID pandemic remains unclear. As the stability of currently available COVID vaccines depends on different temperatures of the “cold chain” for storage and transportation, reliance on these vaccines alone for global vaccination may be difficult. In this context, the stability of the lyophilized smallpox vaccine, from which the v-NY vector was derived, may present certain advantages. Thus, these findings from deployment of our recombinant v-NY-S and MVA-S vaccine candidates in a prime-boost vaccination regimen may represent a very useful approach to tackling SARS-CoV-2 infections globally.

## Materials and Methods

### Cells, viruses, animals and reagents

BSC40 cells were cultured in DMEM supplemented with 10% fetal bovine serum (FBS) (Gibco) and 1% penicillin-streptomycin (PS) (Gibco). BHK21 cells were cultured in RPMI medium supplemented with 10% FBS and 1% PS. HuTK*-*143 cells were cultured in MEM medium supplemented with 10% FBS and 1% PS. The v-NY strain of vaccinia virus was grown on BSC40 or HuTK*-*143 cells as described previously (98, 100, 113–116). The MVA strain of vaccinia virus (VR-1508, ATCC) was grown on BHK21 cells. SARS-CoV-2 TCDC#4 (hCoV-19/Taiwan/4/2020) is a local isolate and it was propagated on Vero-E6 cells. Eight-week-old female C57BL/6 mice (Charles River strain) were purchased from BioLASCO Taiwan Co. Ltd. Eight-week-old male and female Syrian hamsters (*Mesocricetus auratus*) were purchased from the National Laboratory Animal Center, Taiwan. All animal protocols were approved by the Institutional Animal Care and Utilization Committee of Academia Sinica and were conducted in strict accordance with the guidelines on animal use and care of the Taiwan National Research Council’s Guide. All antibodies and reagents are listed in Table 1 below.

**Table 1.**
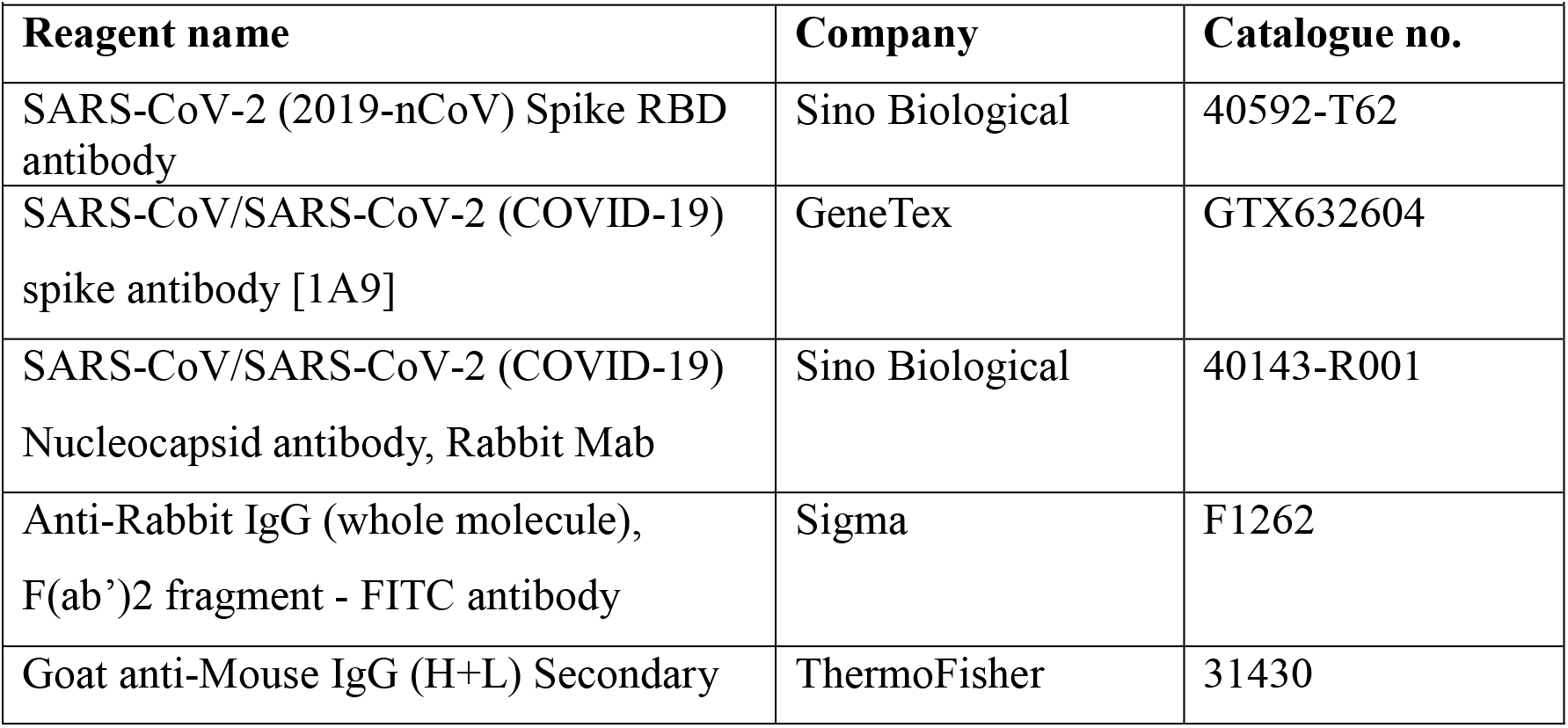
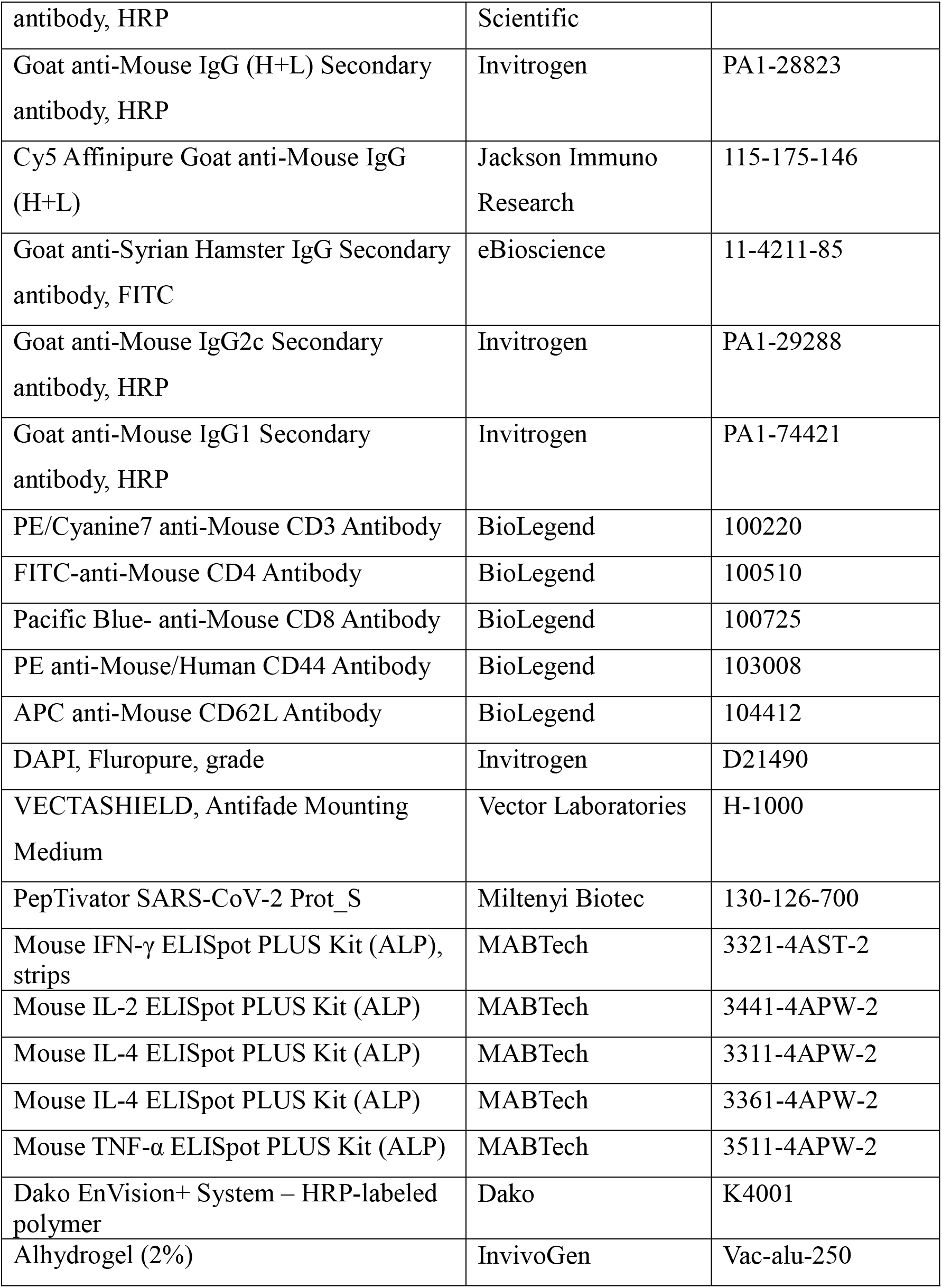

### Construction of recombinant vaccinia viruses

To generate recombinant vaccinia viruses expressing SARS-CoV-2 S protein (Isolate Wuhan-Hu-1, NC_045512), a human codon-optimized open reading frame (ORF) encoding full-length SARS-CoV-2 S protein was inserted into a pSC11 plasmid and under regulatory control by an early and late p7.5k promoter to obtain pSC11-S plasmid (117). The pSC11-S plasmid was transfected into HuTK-143 cells infected with the wild type v-NY virus strain. Lysates were then harvested for multiple rounds of plaque purification of the recombinant virus, named v-NY-S, on HuTK-143 in the presence of 25 μg/ml 5-Bromo-2’-Deoxyuridine (BrdU), as described previously (115). The recombinant MVA strain expressing SARS-CoV-2 S protein, MVA-S, was generated as described for v-NY-S except that BHK21 cells were used and plaque purification was performed in the presence of X-gal (150 μg/ml). Both MVA-S and v-NY-S were subsequently amplified in roller bottles, and the virus stocks were partially purified using a 36% sucrose gradient and titrated prior to use, as described previously (118).

### Immunofluorescence staining of cell surface S protein

BHK21 and BSC40 cells were infected respectively with MVA-S or v-NY-S at a multiplicity of infection (MOI) of 5 PFU/cell for 1 h, washed with PBS, and then incubated in growth media for a further 12 h. The cells were then washed with PBS and fixed with 4% paraformaldehyde, before being immunostained with SARS-CoV- 2 anti-RBD antibody (40592-T62) at a dilution of 1:500 for 1 h at room temperature. Then, the cells were washed with PBS and stained for the secondary antibody FITC-conjugated goat anti-Rabbit IgG Ab (F1262, 1:500 dilution) for 1 h at room temperature, followed by staining with DAPI (5 μg/ml, D21490, Molecular Probes) for 5 min and mounting with Vectashield mounting solution (H-1000, Vector Laboratories). Images were taken using a Zeiss LSM 710 confocal microscope with a 63x objective lens, as described previously (119).

### Prime-boost immunization regimens in mice

Eight-week-old female C57BL/6 mice were housed in the Animal Facility of Academia Sinica (Taipei, Taiwan) for at least 3 days prior to vaccination experiments. We primarily used three regimens of two-dosage prime/boost immunization (depicted in Fig. 2A): (1) MVA5/MVA1 - intramuscular (i.m.) inoculation of MVA-S in the right hind limb at 5×10^7^ PFU/animal, followed by an i.m. boost 4 weeks later of 1×10^7^ PFU/animal of MVA-S virus; (2) vNY1/MVA1 - tail scarification (t.s) of v-NY-S at 1×10^7^ PFU/animal, followed by an i.m. boost 4 weeks later of 1×10^7^ PFU/animal of MVA-S virus into the right hind limb; and (3) vNY5/MVA1- t.s. of v-NY-S at 5×10^7^ PFU/animal, followed by an i.m. boost 4 weeks later of 1×10^7^ PFU/animal of MVA-S virus into right hind limb. Additional prime/boost combinations were designed in order to obtain mouse sera for pseudotyped SARS-CoV-2 virus neutralization assays described in the section below: (4) vNY1/vNY1 - tail scarification (t.s) of v-NY-S at 1×10^7^ PFU/animal, followed by an i.m. boost 4 weeks later of 1×10^7^ PFU/animal of v- NY-S virus into the right hind limb; (5) vNY1/rS - tail scarification (t.s) of v-NY-S at 1×10^7^ PFU/animal, followed by an i.m. boost 4 weeks later of 5 μg recombinant spike protein (aa 14-1209) in 1% alhydrogel (Invivogen) into the right hind limb; and (6) vNY0.1(i.n.)/MVA1- intranasal infection (i.n.) of v-NY-S at 1×10^6^ PFU/animal, followed by an i.m. boost 4 weeks later of 1×10^7^ PFU/animal of MVA-S virus into the right hind limb. As immunization controls, PBS buffer was used as a placebo vaccine for both priming and boosting shots. Blood was collected from immunized mouse cheeks 4 weeks after priming and 2 weeks after boosting, as described previously (120, 121). Sera were prepared from blood and saved at -80 °C until use.

### Prime-boost immunization regimens in Syrian hamsters

Eight-week-old male and female Syrian hamsters were housed in the Animal Facility of Academia Sinica (Taipei, Taiwan) for at least 3 days prior to vaccination experiments. We used three regimens of two-dosage prime/boost immunization (depicted in Fig. 2A): (1) MVA5/MVA1- intramuscular (i.m.) inoculation of MVA-S in the right hind limb at 5×10^7^ PFU/animal, followed by an i.m. boost 4 weeks later of 1×10^7^ PFU/animal of MVA-S virus; (2) vNY1/MVA1- tail scarification (t.s) of v-NY- S at 1×10^7^ PFU/animal, followed by an i.m. boost 4 weeks later of 1×10^7^ PFU/animal of MVA-S virus into the right hind limb; and (3) vNY5/MVA1- t.s. of v-NY-S at 5×10^7^ PFU/animal, followed by an i.m. boost 4 weeks later of 1×10^7^ PFU/animal of MVA-S virus into right hind limb. As immunization controls, PBS buffer was used as a placebo vaccine for both priming and boosting shots. Blood was collected from immunized hamster gingival veins 4 weeks after priming and 2 weeks after boosting, as described previously (120, 121). Sera was prepared from blood and saved at -80 °C until use.

### Immunoblotting

To measure SARS-CoV-2 S protein expression in cells infected with recombinant viruses, BSC40 and BHK21 cells (5×10^5^) were infected with v-NY-S and MVA-S, respectively, at an MOI of 5 PFU/cell and incubated for 12 h prior to cell harvesting. Cells were lysed with sample buffer and proteins were separated by sodium dodecyl sulfate-polyacrylamide gel electrophoresis (SDS-PAGE). Proteins were then transferred to nitrocellulose membranes (BioRad) using a wet transfer apparatus (Bio- Rad). The membranes were blocked in 5% non-fat milk solution at room temperature (r.t.) for 1 h and incubated overnight with SARS-CoV-2 spike S2 mouse mAb (GTX632604, 1:1000 dilution) at 4 °C. The blots were then washed three times with PBST (PBS containing 0.1% Tween-20), incubated at r.t. with HRP goat anti-mouse IgG Ab (31430, 1:20,000) for 1 h and developed using a Western Lightening Enhanced Chemiluminescence kit (PerkinElmer) according to the manufacturer’s protocol.

To test reactivity of immunized mouse and hamster sera to SARS-CoV-2 spike protein, the extracellular domain of spike protein from residue 14 to 1209, consisting of S1 and S2 but without the transmembrane domain, was expressed in HEK 293 cells and subsequently purified. The purified spike protein contained human complex type glycans, and exists as a trimer in solution with an apparent molecular weight between 170 to 235 kDa on SDS-PAGE (monomer), and ∼600 kDa (trimer) on Superose 6 size-exclusion chromatography. Purified spike protein (20 ng/well) was separated by SDS-PAGE, transferred to nitrocellulose membranes and blocked in 5% non-fat milk solution at r.t. as described above. The membrane was separated into multiple strips and each strip was incubated overnight with individual sera collected from immunized mice (1:100 dilution) or hamsters (1:50) at 4 °C. These blots were then washed three times with PBST, incubated at r.t. with HRP goat anti-mouse (31430, 1:20,000) or HRP goat anti-hamster (PA1-28823, 1:5,000) antibodies for 1 h at r.t. and then developed using a Western Lightning Enhanced Chemiluminescence kit (PerkinElmer) according to the manufacturer’s protocol.

### Flow cytometry analysis of cell-surface SARS-CoV-2 S protein expression

To detect spike protein expression on the surface of cells infected with MVA-S or v-NY-S, BSC40 and BHK21 cells (5×10^5^) were infected with v-NY-S and MVA-S, respectively, at an MOI of 5 PFU/cell and incubated for 12 h, before being detached via treatment with 2 mM EDTA in PBS. Cells were incubated with SARS-CoV-2 anti-RBD antibody (40592-T62, 1:500) at 4°C for 1 h. The cells were then washed with FACS buffer (PBS containing 2% FBS), stained with FITC-conjugated goat anti-Rabbit IgG Ab (F1262, 1:500) for 1 h at 4°C, washed with FACS buffer and analyzed by flow cytometry (BD LSR-II, BD Biosciences).

To detect anti-spike antibody in the sera of immunized mice and hamsters, SF9 insect cells were infected with either wild type baculovirus (WT-BAC) or a recombinant baculovirus (S-BAC) that expressed a chimeric SARS-CoV-2 S-gp64 protein in which the transmembrane and C-terminal regions of S protein were replaced by the transmembrane and C-terminal regions of baculovirus GP64 so that the S-gp64 fusion protein would be expressed on insect cell surfaces. These cells were cultured for 48 h before incubating with mouse (1:100 dilution) or hamster (1:20 dilution) serum in FACS buffer for 1 hour on ice. After two washes with FACS buffer, the cells were incubated with Cy5-Goat anti-mouse IgG Ab (115-175-146, 1:500) or FITC-Goat anti-hamster IgG Ab (11-4211-85, 1:100) for 30 min on ice, washed twice, resuspended in FACS buffer containing propidium iodide, and then analyzed by flow cytometry (BD LSR-II).

### SARS-CoV-2 pseudotyped virus neutralization assay

Lentiviral vectors pseudotyped with spike protein of wild type SARS-CoV-2 and SARS-CoV-2 variants of concern (VOC) (Table 2) were generated and titered by the National RNA Technology Platform and Gene Manipulation Core, Academia Sinica, Taipei, Taiwan. Neutralization assays on pseudotyped virus were performed by the same core facility as described previously (122), but with minor modifications. In brief, 1,000 units of the pseudotyped lentivirus with SARS-CoV-2 S protein were incubated at 37 °C for 1 h with serially-diluted sera obtained from vaccinated animals. The mixture was then added to HEK-293T cells expressing human ACE2 receptor (10^4^ cells/well of a 96-well plate) and incubated for 24 h at 37 °C. This cell culture was then replaced with 100 μl of fresh DMEM plus 10% FBS, and the cells were incubated for another 48 h before undergoing luciferase assay. The reciprocal dilution of serum required for 50% inhibition of virus infection (ND_50_) was assessed by measuring luciferase intensity.

**Table 2.**

| **Table 2** |  |  |  |
| --- | --- | --- | --- |
| **Scientific name** | **WHO label\*** | **Mutation(s) in Spike protein (mutations in the RBD are in red)** | **Note** |
| Wild Type (Wuhan-Hu-1 isolate) |  | NA | NC_045512.2 |
| B.1.1.7 | Alpha | 69-70 del, Y144 del, **N501Y**, A570D, D614G, P681H, T716I, S982A, D1118H | VOC |
| B.1.351(501Y.V2) | Beta | L18F, D80A, D215G, 242-244 del, R246I, **K417N**, **E484K**, **N501Y**, D614G, A701V | VOC |
| P.1 | Gamma | L18F, T20N, P26S, D138Y, R190S, **K417T**, **E484K**, **N501Y**, D614G, H655Y, T1027I | VOC |
| B.1.617 |  | G142D, E154K, **L452R**, **E484Q**, D614G, P681R, Q1071H, H1101D | - |
| B.1.617.2 | Delta | T19R, G142D, 156-157 del, R158G, **L452R**, **T478K**, D614G, P681R, D950N | VOC |

\* https://www.who.int/en/activities/tracking-SARS-CoV-2-variants/. RBD- receptor
binding domain; VOC- variant of concern.

### SARS-CoV-2 neutralization assay

Serially diluted antibodies from immunized mice or hamsters were incubated at 37 °C for 1 h with 100 TCID_50_ SARS-CoV-2 TCDC#4 (hCoV-19/Taiwan/4/2020). The mixtures were then added to pre-seeded Vero E6 cells for a 4-day incubation. Cells were fixed with 10% formaldehyde and stained with 0.5% crystal violet for 20 min. The plates were washed with tap water and scored for infection. The 50% protective titer was calculated according to the Reed & Muench Method (123).

### Immunoglobulin ELISA for SARS-CoV-2 S-specific antibodies

Immunoglobulin ELISA was performed as described previously (17) with some modifications. Recombinant SARS-CoV-2 S protein (10 ng/well) was coated onto a 96-well plate (Costar assay plate, Corning, 3369) for 24 h at 4 °C. The plates were then washed with PBST and blocked with 1% BSA in PBS solution for 1 h, followed by washes with PBST. Coated plates were incubated for 1 h at r.t. with sera that had been serially diluted in PBS containing 1% BSA, then washed with PBST, and incubated with HRP-conjugated IgG2C (PA1-29288, 1:15000) or HRP-conjugated IgG1 (PA1-74421, 1:6000) secondary antibodies at r.t. for 1 h. Plates were washed with 5x PBST and incubated with commercial TMB substrate for color development (Clinical Science Products Inc.). To stop the reaction, 2N H_2_SO_4_ was added and the plates were read at an optical density of 450 nm using an ELISA reader. End-point titers were calculated as the serum dilutions that emitted an optical density (O.D) greater than four times the background level (secondary antibody only), as described previously (17).

### ELISpot assay of mouse splenocytes

ELISpot assays to monitor cytokine levels in splenocytes stimulated with a SARS-CoV-2 spike peptide pool were performed essentially as described previously (111, 124). In brief, spleens were collected from immunized mice four weeks after vaccine boosting. We mixed 4×10^5^ splenocytes with a peptide pool of SARS-CoV-2 S protein sequences (Miltenyi Biotech, 130-126-700) at 1 μg/ml concentration in 100 μl medium (RPMI + 10% FBS + 1% PS), and then incubated them for 24 h at 37 °C in ELISpot plates (MABTECH) precoated with IFN-γ (3321-4AST-2), IL-2 (3441- 4APW-2), IL-4 (3311-4APW-2), IL-6 (3361-4APW-2) or TNF-α (3511-4APW-2).

Cells were then washed with 5x PBS and the ELISpots were developed according to the manufacturer’s protocol and quantified using an AID vSpot machine.

### Analyses of T effector memory (Tem) cells

Flow cytometric analyses of Tem cells were performed as described previously (16) with minor modifications. Splenocytes were isolated from immunized mice at 4 weeks after vaccine boosting. After depleting red blood cells with Ammonium-Chloride-Potassium (ACK) lysis buffer, splenocytes were stimulated with 1 μg/ml of a SARS-CoV-2 spike-specific peptide pool (Miltenyi Biotech, 130-126-700) in medium (RPMI + 10% FBS + 1% PS) for 2 h at 37°C. The cells were subsequently washed twice with FACS buffer, and then incubated with an antibody cocktail including anti-CD3-PE/Cyanine7, anti-CD4-FITC, anti-CD8-Pacific blue, anti-CD44- PE and anti-CD62L-APC for 15 min on ice. The cells then underwent fluorescence-activated cell sorting (FACS) analyses, whereby CD4^+^ or CD8^+^ subpopulations were first gated from total splenocytes, and then further gated for CD44^+^CD62L^-^ as Tem cells. Dead cells were stained with eFluor 506 viability dye (eBioscience). Cells were acquired using a BD LSR II (BD Biosciences) flow cytometer and data analyses were performed with FlowJo 8.7 software.

### Syrian hamster challenge experiments

Syrian hamsters were immunized according to one of the three prime-boost vaccination regimens described above, anesthetized, with Zoletil-50 (50mg/kg) and then intranasally (i.n) challenged with 1×10^5^ PFU of SARS-CoV-2 TCDC#4 (hCoV- 19/Taiwan/4/2020, GISAID accession ID: EPI_ISL_411927) (lot: IBMS20200819, 8×10^5^ PFU/ml) in a volume of 125 μl. All animals were weighed daily after SARS-CoV-2 challenge. At 3 and 7 days post infection (d.p.i.), lungs were harvested for SARS-CoV-2 virus titer determination, viral RNA quantification and histopathological examination. Differences in body weight between experimental groups of animals were analyzed statistically using a two-tailed unpaired Student’s *t* test.

### Quantification of viral titers in lung tissues by cell culture infection assay

The middle, inferior, and post-caval lobes of hamsters at 3 and 7 days post challenge with SARS-CoV-2 were homogenized in 4ml of DMEM with 2% FBS and 1% PS using a homogenizer. Tissue homogenate was centrifuged at 15,000 rpm for 5 min and the supernatant was collected for live virus titration. Briefly, 10-fold serial dilutions of each sample were added in quadruplicate onto a Vero E6 cell monolayer and incubated for 4 days. Cells were then fixed with 10% formaldehyde and stained with 0.5% crystal violet for 20 min. The plates were washed with tap water and scored for infection. The fifty-percent tissue culture infectious dose (TCID_50_)/ml was calculated according to the Reed & Muench Method (123).

### Real-time RT-PCR for SARS-CoV-2 RNA quantification

To measure the RNA levels of SARS-CoV-2, specific primers targeting nucleotides 26,141 to 26,253 of the SARS-CoV-2 envelope (E) gene were used for real-time RT-PCR, as described previously (125), forward primer E-Sarbeco-F1 (5’-ACAGGTACGTTAATAGTTAATAGCGT-3’), reverse primer E-Sarbeco-R2 (5’- ATATTGCAGCAGTACGCACACA-3’), probe E-Sarbeco-P1 (5’-FAM-ACACTAGCCATCCTTACTGCGCTTCG-BBQ-3’). A total of 30 μl RNA solution was collected from each sample using an RNeasy Mini Kit (QIAGEN, Germany) according to the manufacturer’s instructions. RNA sample (5 μl) was added into a total 25-μl mixture of the Superscript III one-step RT-PCR system with Platinum Taq Polymerase (Thermo Fisher Scientific, USA). The final reaction mix contained 400 nM of the forward and reverse primers, 200 nM probe, 1.6 mM deoxy-ribonucleoside triphosphate (dNTP), 4 mM magnesium sulfate, 50 nM ROX reference dye, and 1 μl of the enzyme mixture. Cycling conditions were performed using a one-step PCR protocol: 55°C for 10 min for first-strand cDNA synthesis, followed by 3 min at 94°C and 45 amplification cycles at 94°C for 15 sec and 58°C for 30 sec. Data was assessed using an Applied Biosystems 7500 Real-Time PCR System (Thermo Fisher Scientific). A synthetic 113-basepair oligonucleotide fragment was used as a qPCR standard to estimate copy numbers of the viral genome. The oligonucleotides were synthesized by Genomics BioSci & Tech Co. Ltd. (Taipei, Taiwan).

### Histopathology

The left lung of each hamster at 3 and 7 days post challenge with SARS-CoV-2 was removed and fixed in 4% paraformaldehyde for 1 week. The lung samples were then embedded, sectioned, and stained with Hematoxylin and Eosin (H&E), followed by microscopic examination. Immunohistochemical staining was performed with a monoclonal rabbit anti-SARS-CoV/SARS-CoV-2 nucleocapsid (NP) antibody (1:1000, 40143-R001, Sino Biological), followed by incubation with Dako EnVision+ System HRP. Brownish signals were subsequently developed upon addition of 3,3’ diaminobenzidine (DAB) and counterstained with hematoxylin. Images were photographed using a Zeiss Axioimager-Z1 microscope with 4x and 20x objective lenses.

### Statistical analyses

Statistical analyses were conducted using Student’s *t* test in Prism (version 9) software (GraphPad). For multiple comparisons, the p values were adjusted by the “fdr” method using the “p.adjust” function in R v4.0.4. Adjusted p values <0.05 were considered statistically significant. *p<0.05; **p<0.01; ***p<0.001; ****p<0.0001.

## Acknowledgements

We thank Sue-Ping Lee of the Imaging Core Facility, Ya-Min Lin of the FACS Core Facility and Kun-Hai Yeh of Bioinformatics Core facility at the Institute of Molecular Biology and Data Science Statistical Cooperation Center-Statistical Clinics at Acadmia Sinica. We also thank Brad Cleveland for advice and assistance with the v-NY vector.

## Supporting information captions

**S1 Fig.** Weight change in C57BL/6 mice after immunization with one of three regimens.

**S2 Fig.** (A). Images of skin scarification in Syrian hamsters at days 5,10 and 15 after primary immunization. (B) Weight change in Syrian hamsters after immunization with one of the three regimens.

**S3 Fig.** TCID_50_ value of SARS-CoV-2 in lung tissues of hamsters at 7 d.p.i after SARS-CoV-2 challenge: PBS/PBS control (n=3); MVA5/MVA1 (n=5), The dotted line represents assay limits of detection.

