## Supplementary figures and images for "Vaccinia virus-based vaccines confer protective immunity against SARS-CoV-2 virus in Syrian hamsters"

### Supplemental Figure 1

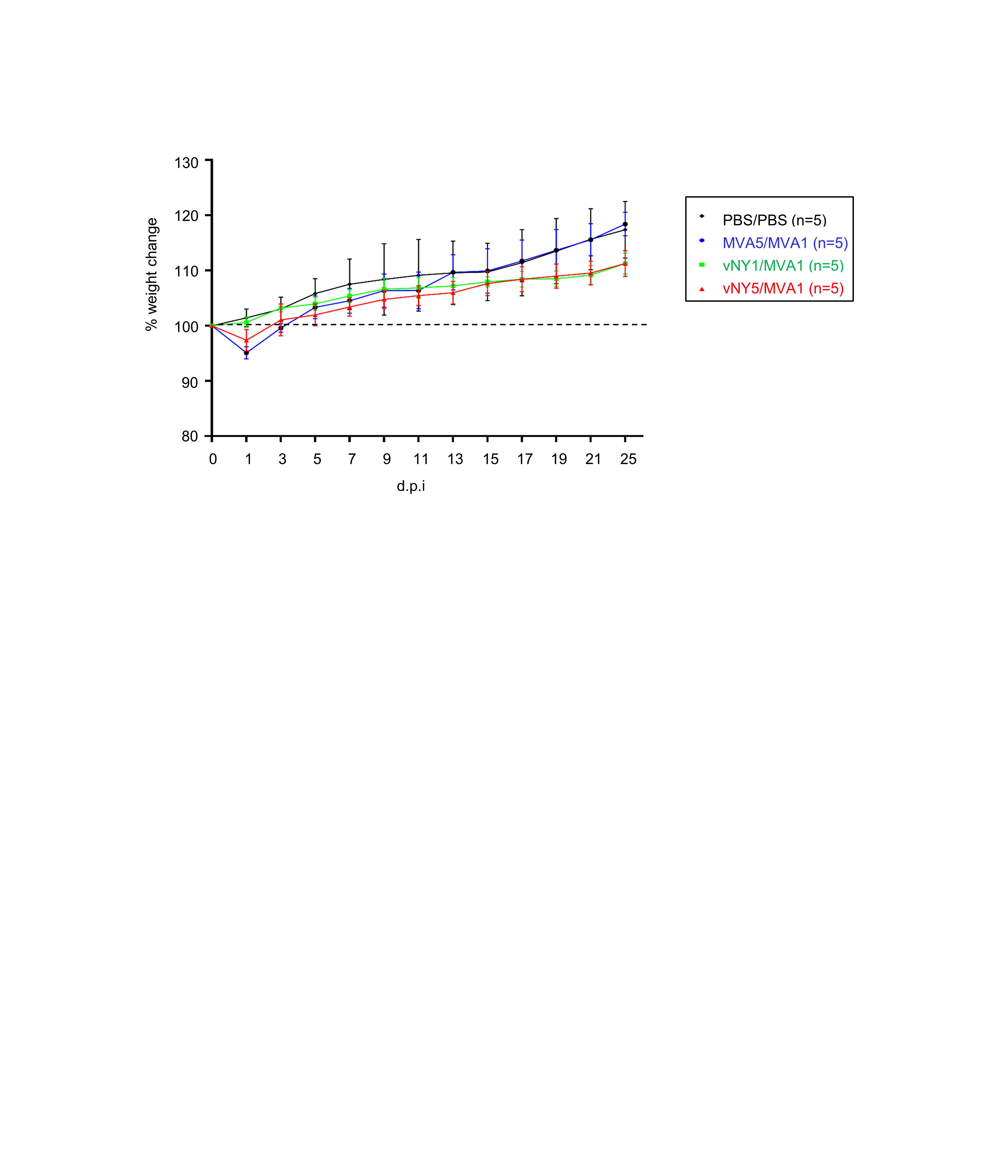

### Supplemental Figure 2

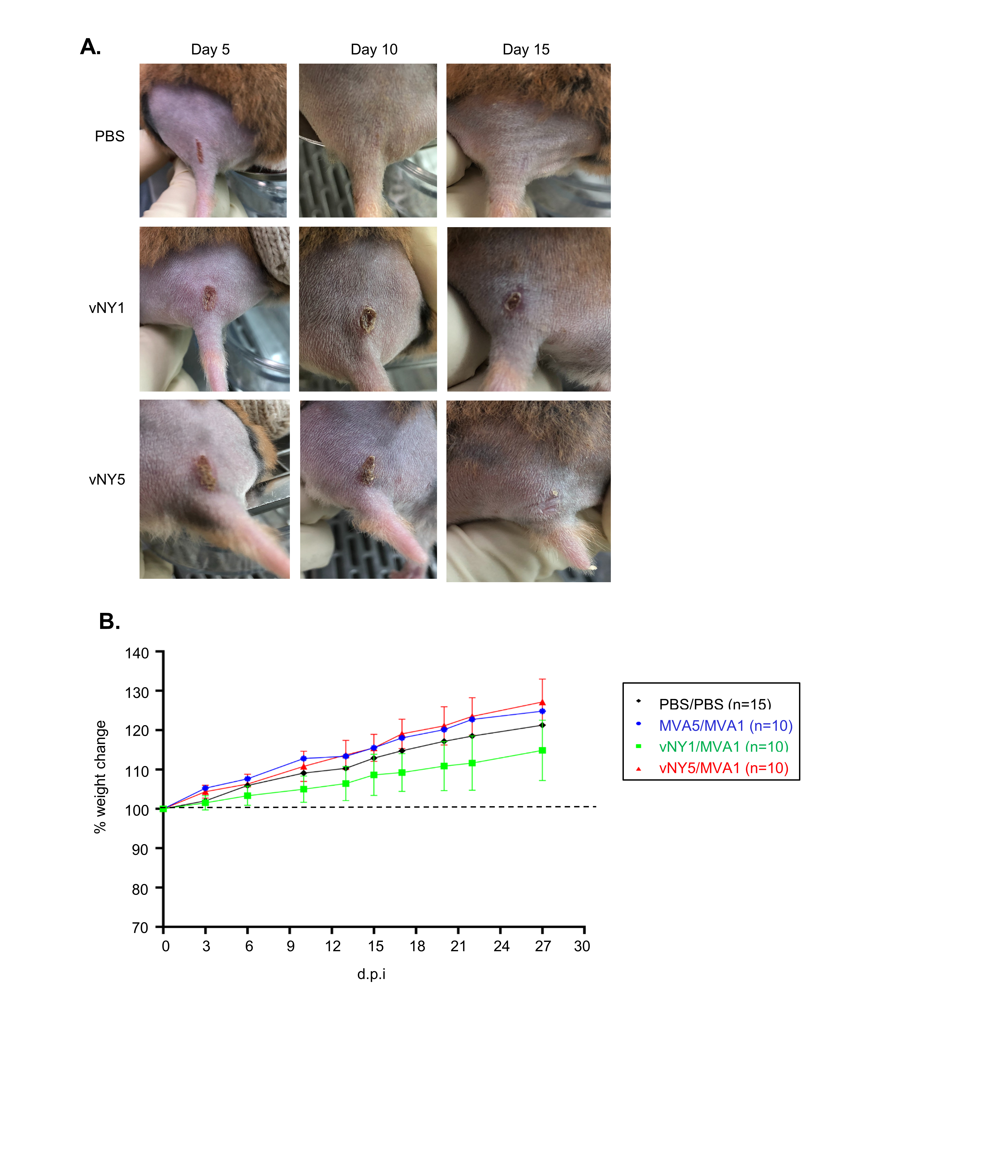

### Supplemental Figure 3

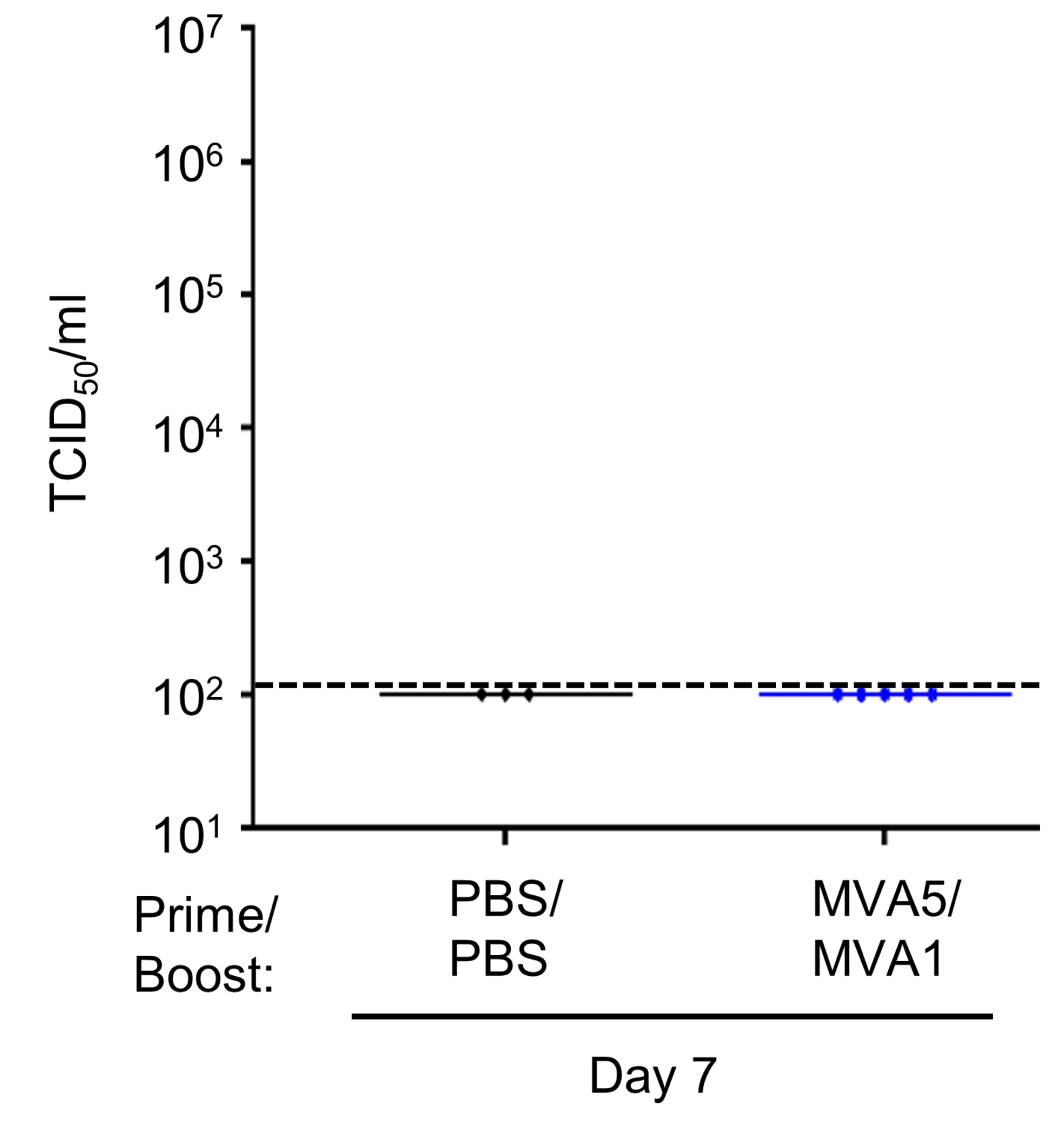
